# A human immune/muscle xenograft model of FSHD muscle pathology

**DOI:** 10.1101/2023.11.17.567590

**Authors:** Katelyn Daman, Jing Yan, Lisa M. Burzenski, Jamie Kady, Leonard D. Shultz, Michael A. Brehm, Charles P. Emerson

## Abstract

**Background:** Facioscapulohumeral muscular dystrophy (FSHD) disease progression is associated with muscle inflammation, although its role in FSHD muscle pathology is unknown.

**Methods:** We have developed a novel humanized mouse strain, NSG-SGM3-W41, that supports the co-engraftment of human hematopoietic stem cells (HSCs) and muscle myoblasts as an experimental model to investigate the role of innate immunity in FSHD muscle pathology.

**Results:** The NSG-SGM3-W41 mouse supports the selective expansion of human innate immune cell lineages following engraftment of human HSCs and the co-engraftment and differentiation of patient-derived FSHD or control muscle myoblasts. Immunohistological and NanoString RNA expression assays establish that muscle xenografts from three FSHD subjects were immunogenic compared to those from unaffected first-degree relatives. FSHD muscle xenografts preferentially accumulated human macrophages and B cells and expressed early complement genes of the classical and alternative pathways including complement factor C3 protein, which is a mediator of early complement function through opsonization to mark damaged cells for macrophage engulfment. FSHD muscle xenografts also underwent immune donor dependent muscle turnover as assayed by human spectrin β1 immunostaining of muscle fibers and by NanoString RNA expression assays of muscle differentiation genes.

**Conclusions:** The NSG-SGM3-W41 mouse provides an experimental model to investigate the role of innate immunity and complement in FSHD muscle pathology and to develop FSHD therapeutics targeting DUX4 and the innate immunity inflammatory responses.

## Background

Facioscapulohumeral muscular dystrophy (FSHD) is a prevalent epigenetic disease caused by genetic disruptions including contraction in the D4Z4 repeats at the 4qA locus (2, 3) or loss of function mutations in chromatin modifier genes (4, 5) that lead to hypomethylation of the D4Z4 locus and misexpression of the germline transcription factor gene *DUX4,* the FSHD disease gene. DUX4 is encoded by the terminal D4Z4 repeat (6, 7) and normally functions during early germline development (8, 9). DUX4 misexpression in muscle disrupts the muscle transcriptome through activation of a large set of germline-specific genes (10–13) which are surrogate biomarkers of DUX4 expression, although their role in DUX4 muscle pathology is unknown.

DUX4 misexpression alone does not appear to be sufficient to account for FSHD muscle pathology, although the variation in levels of DUX4 could potentially be related to disease severity. While *DUX4* misexpression causes muscle toxicity *in vitro* in patient muscle cells (14) and *in vivo* in inducible mouse models (15, 16), the onset of the muscle pathology in FSHD patients is highly variable including early onset and non-manifesting disease (17). Clinical disease initiates sporadically in facial, scapular, and humeral muscles, which are often affected asymmetrically, and over time involves most muscle groups (18–20). Finally, transient expression of DUX4 in transgenic zebrafish and mouse models results in delayed muscle pathology (21, 22), and transient DUX4 induction in a human myoblast culture model generates H3.X and H3.Y histones marking the DUX4 target genes for delayed muscle toxicity (23).

Our investigations have focused on the role of innate immunity as a modifier and amplifier of FSHD muscle pathology and disease progression. FSHD disease progression includes immune cell infiltration (24, 25) followed by muscle turnover and fatty and fibrotic replacement (12, 20, 21, 26, 27), as evidenced by transcriptome (18, 27, 28) and immunohistology assays of patient muscle biopsies (29) and muscles of DUX4 inducible mouse models (16, 30). DUX4 misexpression inhibits nonsense mediated decay (NMD) (31), leading to the production of abnormally spliced RNAs and aberrant proteins, predicted to elicit Damage Associated Molecular Patterns (DAMPs) known to stimulate an innate immune response (32). A role for innate immunity in FSHD muscle pathology is also supported by high expression level of complement genes in muscle biopsies (20, 28) and increased levels of complement C3 in blood of FSHD patients (33).

To investigate the role of innate immunity in FSHD muscle pathology, we developed a novel humanized HSC/muscle engrafted mouse model, in NSG-SGM3-W41 mice. This mouse strain has been engineered to selectively expand human innate immune cell lineages following engraftment of umbilical cord blood (UCB) derived hematopoietic stem cells (HSCs) in the absence of irradiation preconditioning and supports co-engraftment of patient derived FSHD (or unaffected control) muscle myoblasts into the mouse tibialis anterior (TA) muscle to produce differentiated FSHD human muscle that expresses the FSHD disease gene, *DUX4* (34). Our findings show that FSHD muscle xenografts in HSC engrafted NSG-SGM3-W41 mice preferentially accumulate human macrophages and B cells, express early complement RNAs encoding activators of both the classical and alternative pathways, and upregulate C3 RNA and protein, the mediator of early complement response (35). FSHD muscle xenografts also undergo muscle turnover dependent on specific HSC immune donors, supporting the idea that innate immunity directly contributes to FSHD pathology. A role for late complement in muscle turnover is excluded by our results that FSHD muscle xenografts do not express RNAs encoding the complement membrane attack complex (MAC) (36). Based on our findings, we hypothesize that C3 complement activated by the early complement pathway response to FSHD muscle which produces DAMPs and promotes muscle turnover through opsonization of FSHD muscle for macrophage recognition and engulfment (35).

## Methods

### Mouse model generation

NOD.Cg-*Kit^em1Mvw^Prkdc^scid^IL2rg^tm1Wjl^* Tg(CMV-IL3,CSF2,KITL)Eav/MloySzJ (NSG-SGM3-W41) mice were developed by LDS at the Jackson Laboratory. NSG-SGM3 mice were originally obtained from The Jackson Laboratory. The W41 mutation in the mouse *Kit* gene, consisting of a G to A point mutation in the kinase domain (V831M), was made directly in NSG zygotes using CRISPR-Cas9 and oligo-mediated homology directed repair, as described previously (37). To reduce the potential for off-target mutations, a truncated guide (38) was used to target the sequence: GCACGACTGCCCGTGAAG and the NSG-*Kit^W41^* allele was generated using the donor oligonucleotide template: AGGGGAGGTGGCTGGAGGTCACAAGGTTTAAGGTCCTCGTCTATCGCTGTCTTCATTAGCTGCTTGAATTTGCTGTGTTCCGTTCTAGGCACGACTGCCCATGAAGTGGATGGCACCAGAGAGCATTTTCAGCTGCGTGTACACATTTGAAAGTGATGTCTGGTCCTATGGGATTTTCCTCTGGGAGCTCTTCTCCTTAG. NSG-SGM3 mice were intercrossed with NSG- *Kit^W41^*mice and further crosses were made to fix all genes to homozygosity in NSG-SGM3 *Kit^W41^* mice.

### Isolation of human umbilical cord blood (UCB)-HSC and engraftment into mice

Human UCB was obtained in accordance with the Committee for the Protection of Human Subjects in Research guidelines of the University of Massachusetts Chan Medical School. UCB was provided by the University of Massachusetts Memorial Umbilical Cord Blood Donation Program. Groups of 4- to 8-week-old male and female NSG-SGM3-W41 mice were injected IV with CD3-depleted (Miltenyi Biotech) human UCB containing 1×10^5^ CD34^+^ HSCs (39, 40). At the indicated time points, flow cytometry analyses of the blood of engrafted mice quantified the engraftment of the human immune system. For experimental studies, mice with >10% peripheral human CD45+ cells and >5% human CD3+ T cells were used.

### Flow Cytometry

For analysis of human immune system development in HSC engrafted NSG-SGM3-W41 mice, the following monoclonal antibodies specific for human antigens were used: human CD45 (2D1), CD3 (UCHT1), CD20 (2H7) and CD33 (WM53). Anti-mouse CD45 (30F-11) was used to exclude mouse leukocytes. The antibodies were purchased from BD Biosciences, Inc. (CA) or BioLegend (CA). Single-cell suspensions of the spleens were prepared from engrafted mice, and whole blood was collected in heparin. Single cell suspensions of 5×10^5^ splenic cells in 50 μl or 100 μl of whole blood was washed with FACS buffer (PBS supplemented with 2% fetal bovine serum, (HyClone, UT) and 0.02% sodium azide (Sigma, MO) and then pre-incubated with rat anti-mouse FcR11b (clone 2.4G2, BD Biosciences, CA) to block Fc binding. Specific antibodies against cell surface antigens were then added to the samples and incubated for 30 min at 4°C. Stained samples were then washed and fixed with 2% paraformaldehyde for cell suspensions or treated with BD FACS lysing solution for whole blood. At least 100,000 events were acquired on LSRII instrument (BD Biosciences, CA) or Aurora (Cytek Biosciences, CA). Data analysis was performed with FlowJo software (Tree Star, Inc., OR).

### Cell Culture

CD56+ FACS enriched FSHD and control myoblasts from families 12, 15 and 17 were cultured on 0.1% gelatin (sigma G9391) coated 15 cm dishes in HMP medium (Ham’s F10 (Cellgro 10- 070-CV) supplemented with 20% FBS (Hyclone SH30071.03), and 1% chick embryo extract (made in house)) and passaged using TrypLE (ThermoFisher) when 70% confluence was reached.

### Muscle Xenografts

NSG-SGM3-W41 mice were used in accordance with the Institutional Animal Care and Use Committee (IACUC) at the UMass Chan Medical School. Mice were anesthetized with ketamine/xylazine and their hindlimbs were subjected to 18 Gy of irradiation using a Faxitron CellRad X-ray cabinet (Faxitron Bioptics LLC) to ablate the host mouse satellite cell population. Lead shields were used to limit radiation exposure to hindlimbs only. One day after irradiation, mice were anesthetized using an isoflurane vaporizer (SurgiVet model 100) and Tibialis Anterior (TA) muscles were injected with 50 μl of 1.2% Barium Chloride (Sigma) bilaterally to degenerate mouse muscle. Three days after muscle injury, 1 x 10^6^ CD56+ biopsy- derived myoblasts were resuspended in 50 μl 1 mg/mL laminin (Sigma, L2020) in phosphate buffered saline (PBS) and injected bilaterally into the body of TA muscles. Xenoengrafted mice were euthanized 3-4 weeks post engraftment by CO_2_ asphyxiation followed by cervical dislocation. For immunohistology experiments, TA muscles were embedded in Tissue-Tek O.C.T compound (Sakura) and frozen on liquid nitrogen cooled isopentane and kept at -80°C until cryosectioning. For RNA isolation, xenoengrafted TA muscles were snap frozen in liquid nitrogen and kept at -80°C until RNA isolation.

### RNA isolation for NanoString

RNA was isolated from xenoengrafted TA muscles using Aurum Total RNA Fatty and Fibrous Tissue kit (Bio-Rad) per the manufacturer’s specifications. For NanoString digital RNA quantification, 150 ng of total RNA was used for each xenografted TA muscle. A custom inflammation NanoString panel with human-specific probes for muscle protein genes (*MEF2C*, *MYH8* and *MYL2*), DUX4 target genes (*LEUTX* and *MBD3L2*), inflammation genes and multiple housekeeping genes was used for all analysis on an nCounter Sprint profiler (NanoString Technologies, Seattle, WA). Raw mRNA counts for each TA sample were normalized to a panel of housekeeping genes (*RPL13A*, *GAPDH*, *GUSB, HRPT1*, *PGK1*, *TUBB*, and *VCP*) using nSolver software (NanoString Technologies, Seattle, WA).

### TA sectioning and immunohistology

Frozen TA muscles embedded in Tissue-Tek OCT compound (Sakura) were cryosectioned using a Leica CM3050 S Cryostat. Tissue sections 10 μm thick were mounted onto Superfrost Plus glass microscope slides (Fisher Scientific) and kept at -20°C. When thawed, the sections were fixed with ice cold acetone for 10 minutes at -20°C. For lamin A/C (mab636) and spectrin β1 (NCL-SPEC1) co-staining, we employed the “mouse-on-mouse” (M.O.M) kit (Vector Laboratories) to reduce non-specific antibody staining per the manufacturer’s specifications. Antibodies were used sequentially then slides were incubated with Hoechst block for 10 minutes. For human CD45 (Dako, M0701), CD19 (Abcam, ab134114), CD68 (Agilent, clone PG-M1) or human specific C3 (ThermoFisher, JF10-30) immunostaining, primary antibodies were incubated with slides overnight at 4°C followed by 2 x 5 min PBS washes. The corresponding secondary antibodies were added and incubated at room temperature for one hour followed by 2 x 5 min PBS washes. Slides were incubated with Hoechst for 10 mins at room temperature then dried and coverslips were mounted with Fluorogel. Fluorescent images were taken using a Leica DMR fluorescence microscope with IKona monochrome high sensitivity 6MP camera with Sony sensor.

### Statistics

NanoString and immunostaining quantification data are shown as the mean ± SEM. Statistical differences for NanoString RNA expression data and immunofluorescence quantification were evaluated using Welch’s t test and were considered significant when the P value was less than 0.05 (* = P < 0.05, ** = P < 0.01, *** = P < 0.001, **** = P < 0.0001). For flow cytometry data in figure 1 comparing more than 2 means, a 2-way ANOVA was performed followed by Bonferroni’s multiple comparisons test. Statistical analyses were performed using Prism V9 (Graphpad Software LLC).

**Figure 1.**
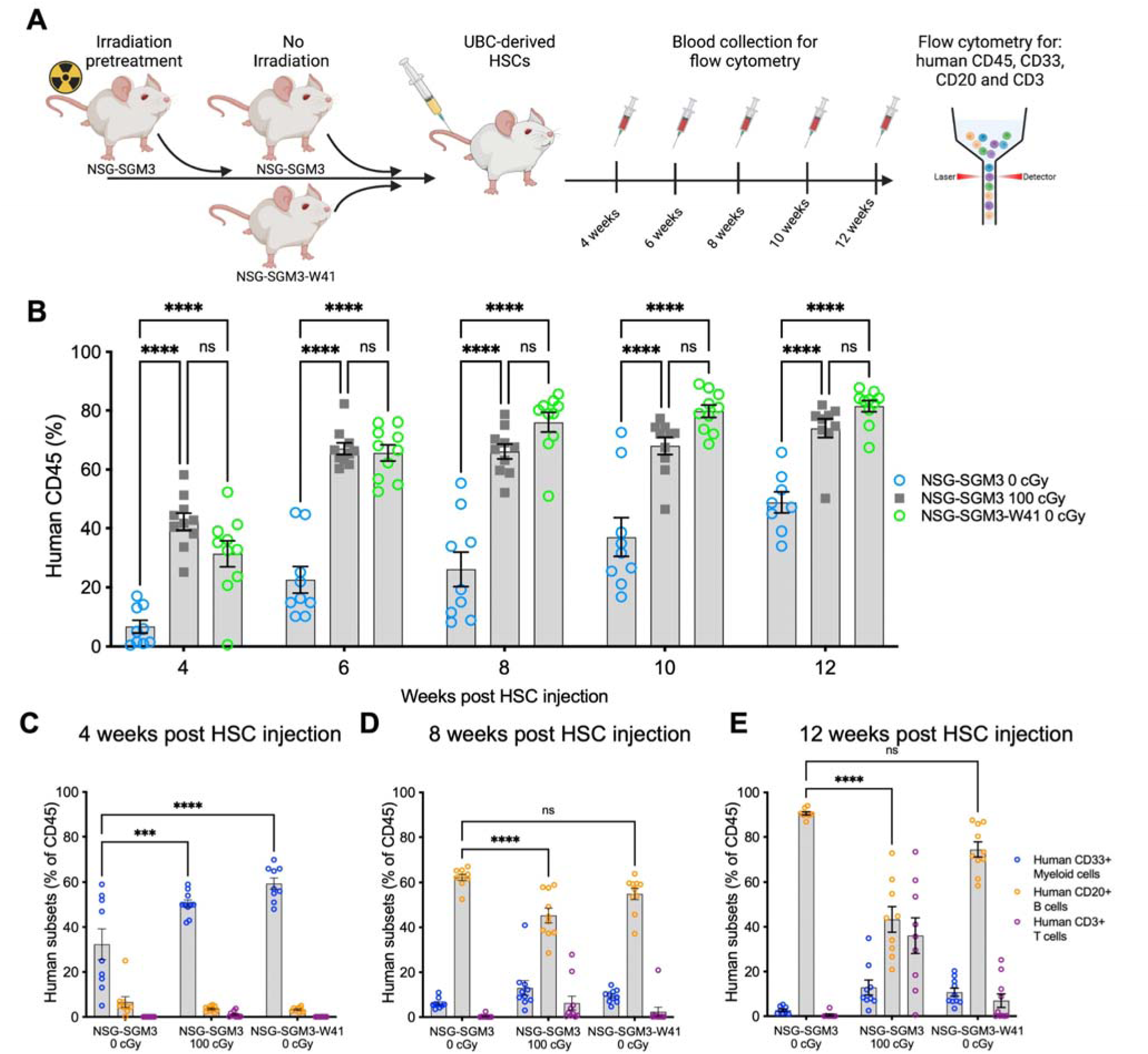
NSG-SGM3-W41 mice support human HSC engraftment and selective expansion of innate immune cells. **(A)** Schematic showing the experimental design comparing the innate immune engraftment of NSG-SGM3 mice with or without 100 cGy irradiation to NSG-SGM3-W41 mice. **(B)** Flow sorting assays of human CD45+ cells in blood at the indicated time points post HSC injection. **(C-E)** Flow sorting assays of the % of CD45+ cells that co-express CD33+ Myeloid cell marker, CD20+ B cell marker, and CD3+ T cell marker in blood at **(C)** 4 weeks, **(D)** 8 weeks or **(E)** 12 weeks post HSC injection. ****=p<0.0001 by 2-way ANOVA with Bonferroni’s multiple comparisons test.

## Results

### Development of the NSG-SGM3-W41 mouse model of human innate immunity

To investigate the role of the human innate immune system in FSHD muscle pathology, we developed a mouse strain (NSG-SGM3-W41) that supports the co-engraftment and differentiation of human CD34+ hematopoietic stem cells (HSCs) and human muscle myoblasts isolated from FSHD and control patient muscle biopsies. The NSG-SGM3-W41 mouse strain was generated first by crossing immune deficient NSG-SGM3 mice that express the human interleukin-3 gene (*IL-3*), human granulocyte/macrophage-colony stimulating factor gene (*GM-CSF*) and human stem cell factor (*SCF*) (41, 42) with NSG mice expressing CRISPR Cas9 W41J point mutation of the *Kit* locus to enable efficient multi-lineage engraftment of HSCs without irradiation (43, 44).

Immune cell development was compared in HSC engrafted NSG-SGM3 mice with or without 100 cGy irradiation pretreatment, and in non-irradiated NSG-SGM3-W41 mice. Mice were injected with 10^5^ CD34+ HSCs isolated from healthy donor UCB, and blood samples from engrafted mice were assayed for circulating human CD45+ immune cells by flow cytometry at 4-, 6-, 8-, 10- and 12-weeks post HSC engraftment (Figure 1A; Additional file 1). Results were expressed as the percentages of human CD45+ blood cells (Figure 1B). Irradiated NSG-SGM3 mice and unirradiated NSG-SGM3-W41 mice both showed 30-40% circulating human CD45+ hematopoietic cells by 4 weeks post HSC injection and this percentage increased to 70-80% at 12 weeks (Figure 1B). By contrast, the non-irradiated NSG-SGM3 mice injected with HSC had only 6% human CD45+ cells at 4 weeks post HSC engraftment and these cell numbers increased to 48% at 12 weeks (Figure 1B), with significantly lower engraftment efficiency than the irradiated NSG-SGM3 or non-irradiated NSG-SGM3-W41 mice at all time points.

Specific lineages of human immune cells produced in HSC engrafted NSG-SGM3 mice with or without 100 cGy irradiation pretreatment and in non-irradiated NSG-SGM3-W41 mice were compared in the blood using flow cytometry to determine the percent CD45+ cells that expressed CD33, a myeloid cell marker, CD20, a B cell marker, or CD3, a T cell marker. Flow cytometry assays were performed at 4-, 8- and 12-weeks post HSC engraftment (Figure 1C- E). At 4 weeks, mice in all three groups were populated primarily by CD33+ myeloid cells, with unirradiated NSG-SGM3 having the lowest engraftment levels (a mean of 32%) compared to the irradiated NSG-SGM3 mice (50%), and NSG-SGM3-W41 mice (59%) (Figure 1C), whereas CD20+ B cell numbers were very low and CD3+ T cells were absent. At 8 weeks post HSC injection, all three groups had lower levels of CD33+ myeloid cells compared to 40 and 60% of CD20+ B cells, and several animals in the irradiated NSG-SGM3 group and one in the

NSG-SGM3-W41 group had a small number of CD3+ T cells (Figure 1D). At 12 weeks post HSC injection, the blood of irradiated NSG-SGM3 mice had robust engraftment of CD33+ myeloid cells, CD20+ B cells and CD3+ T cells; non-irradiated NSG-SGM3 mice primarily showed CD20+ B cell engraftment; and NSG-SGM3-W41 mice showed robust engraftment of CD20+ B cells, moderate levels of CD33+ myeloid cells and low levels of CD3+ T cells (Figure 1E). These data show that NSG-SGM3-W41 mice at 8-weeks post HSC engraftment supported robust development of myeloid and B cells and restricted development of T cells, providing a model for HSC co-engraftment with muscle myoblasts to investigate innate immune responses to FSHD muscle.

### Engraftment of FSHD and control muscle myoblasts in HSC engrafted NSG-SGM3-W41 mice

We next investigated whether HSC engrafted NSG-SGM3-W41 mice would support the engraftment and differentiation of patient FSHD and unaffected control muscle myoblasts. NSG-SGM3-W41 mice were engrafted with 10^5^ CD34+ HSCs from healthy UCB donors at 4 weeks. Two to three weeks after HSC engraftment, the hindlimbs of the HSC engrafted mice were irradiated to block the growth of host mouse muscle satellite cells and Tibialis Anterior (TA) muscles were then injured by barium chloride injection to destroy mouse TA muscle fibers and create a niche for the engraftment and differentiation of human muscle-biopsy derived myoblasts (Figure 2A). Barium chloride injured TA muscles were then engrafted with 10^6^ CD56+ myoblasts isolated from muscle biopsies from three FSHD families (12, 15 and 17), including affected FSHD patients (12A, 15A and 17A) and unaffected control first-degree relatives (12U, 15V and 17U) (Figure 2B)(34). High (17A), medium (12A), and low (15A) DUX4 expressing FSHD cell lines were chosen for experiments to characterize the immune response to FSHD muscle (Figure 2B)(1). Family 12 muscle myoblasts (12A/12U) were co-engrafted into TA muscles of mice with one HSC donor. Family 15 (15A/15V) muscle myoblasts were co- engrafted with two different HSC donors (15D1 and 15D2), and family 17 muscle myoblasts were engrafted into TA muscles of mice engrafted with HSCs from four different donors (17D1, 17D2, 17D3 and 17D4) (Figure 2C). Three to four weeks after muscle stem cell engraftment, the mice were euthanized, their spleens were isolated to evaluate the development of human B cells, myeloid cells and T cells, and their TA muscles were processed for immunohistology or RNA expression analyses (Figure 2A).

**Figure 2.**
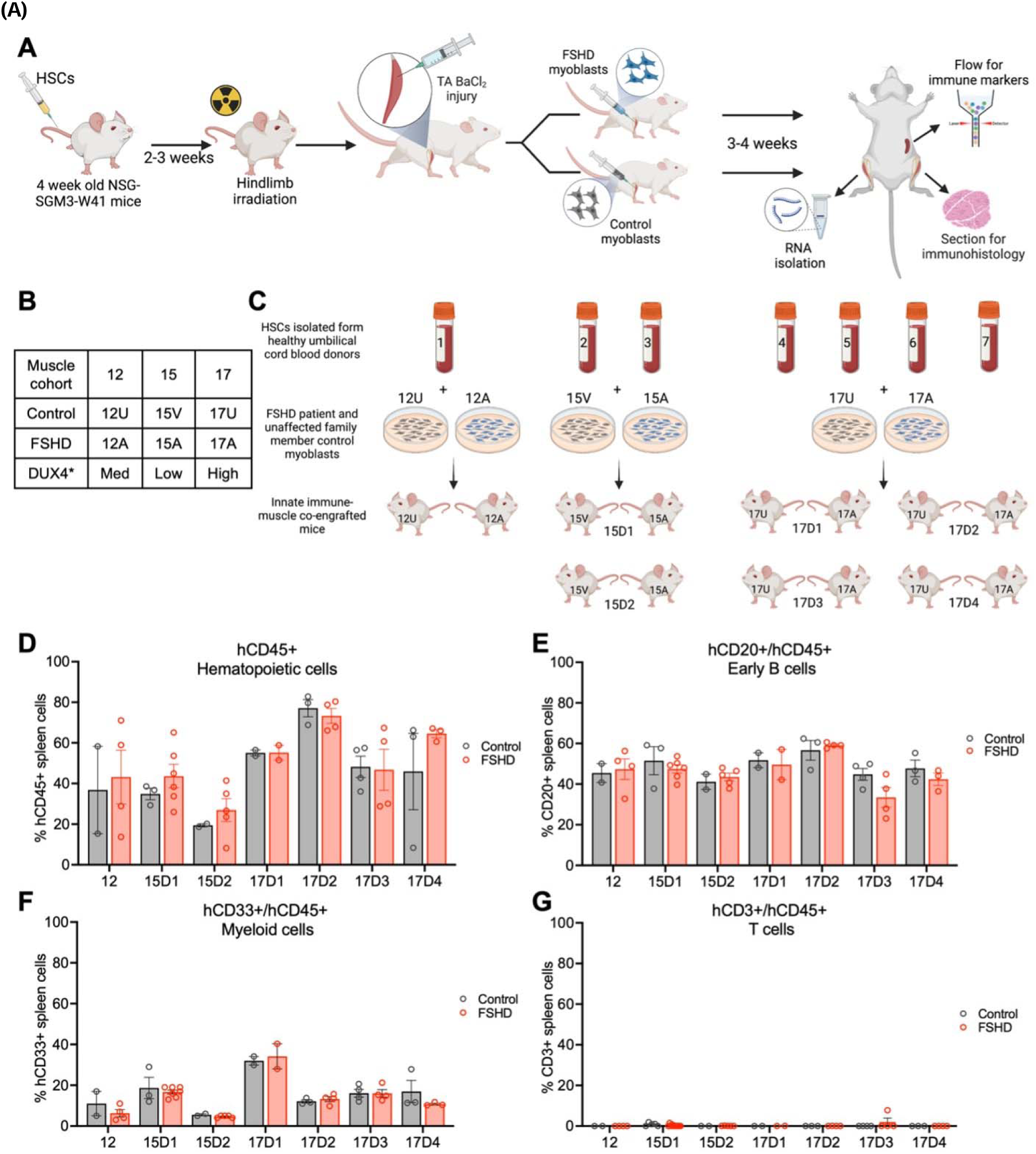
Co-xenoengraftment of human innate immune cells and skeletal muscle in NSG-SGM3-W41 mice. **(A)** Schematic showing the co-engraftment protocol for human donor HSCs and FSHD and control muscle myoblasts and processing of TA muscle xenografts for flow cytometry to identify cell lineages, RNA isolation for qPCR and NanoString, and sectioning for immunohistology. **(B)** Table describing myoblast cell lines used. DUX4 expression levels previously described (1). **(C)** Schematic describing HSC and muscle experimental groups for six HSC donors and three FSHD family cohorts. **(D-G)** Flow cytometry assays the percent hCD45+ cells in spleen **(D**), hCD45/CD20+ B cells **(E),** hCD45/CD33+ Myeloid cells **(F)** or hCD45/CD3+ T cells **(G)** from the indicated engraftment conditions. Each dot represents data from one mouse with bars showing mean per condition ± SEM. Schematics in A and C were

HSC immune engraftment was assessed by flow cytometry analysis of spleen cells assayed for CD45+ hematopoietic lineage cells, CD45+/CD20+ B cells, CD45+/CD33+ myeloid cells, and CD45+/CD3+ T cells. Mice engrafted with all combinations of HSC donors and muscle stem cell donors produced comparable levels of hematopoietic CD45+ cells (Figure 2D), B cells (Figure 2E), and myeloid cells (Figure 2F). A representative flow cytometry gating strategy is shown in Additional file 1A. None of the HSC engrafted mice produced CD45+/CD3+ T cells, confirming that NSG-SGM3-W41 mice are permissive for the expansion of innate immune cell lineages but not T cell lineages during an 8-week period following HSC engraftment (Figure 2G). A minority of mice did not engraft with HSCs based on undetectable CD45+ staining by flow cytometry of spleen cells (Additional file 1B) and by immunohistology of TA muscles (Additional file 1C).

### Enhanced accumulation of CD45+ innate immune cells in FSHD xenograft muscle compared to control muscle

To investigate whether human innate immune cells preferentially infiltrate FSHD muscle xenografts, TA muscles were immunostained for human specific CD45 to identify HSC derived innate immune cells and Hoechst to identify all nuclei (Figure 3A). While CD45+ immune cells were identified in both FSHD and control TA xenografts (Figure 3A), CD45+ cells were significantly more abundant in FSHD than control muscle xenografts in 6 of the 7 immune donors (Figure 3B), reflecting either increased infiltration and/or expansion of human immune cells in FSHD xenografts. To determine whether the human CD45+ cells localized to engrafted human muscle, serial sections of TA muscles were immunostained for human CD45 to show human leukocytes or human spectrin β1 to show human muscle fibers. CD45+ cells colocalized with spectrin β1+ muscle fibers in FSHD xenografts compared to control muscle (Figure 4C), providing evidence that immune cells are trophic to FSHD muscle.

**Figure 3.**
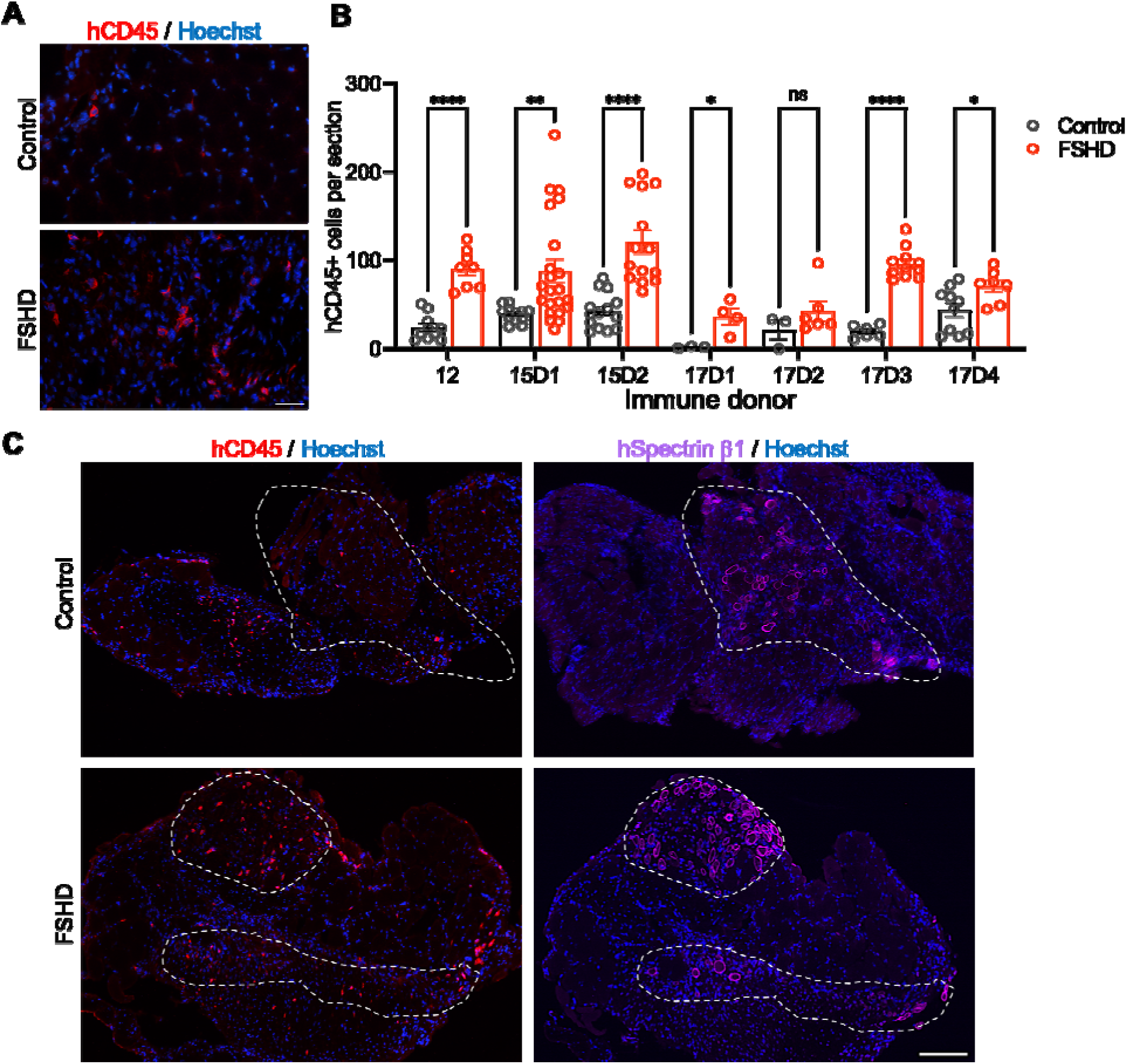
Enhanced accumulation of human CD45+ innate immune cells in FSHD muscle xenografts. **(A)** Humanized TA muscles were cryosectioned and immunostained with human-specific anti-CD45 to identify hematopoietic cells and Hoechst to identify nuclei. Representative images of FSHD and Control engrafted TA muscles are shown. Scale bar = 50 µm. **(B)** Quantification of CD45+ cells per TA muscle section for the indicated engraftment conditions. Each dot shows the number of human CD45+ cells in one muscle section with bars showing mean ± SEM per condition. *=p<0.05, **=p<0.01, ***=p<0.001, ****=p<0.0001 by Welch’s t-test. **(C)** Serial sections were immunostained with human-specific anti-CD4 to identify immune cells or spectrin β1to identify human fibers and Hoechst and nuclei. The humanized muscle are is highlighted in white based on the localization of spectrin β1 muscle fibers and is transposed onto the CD45 immunostained

**Figure 4.**
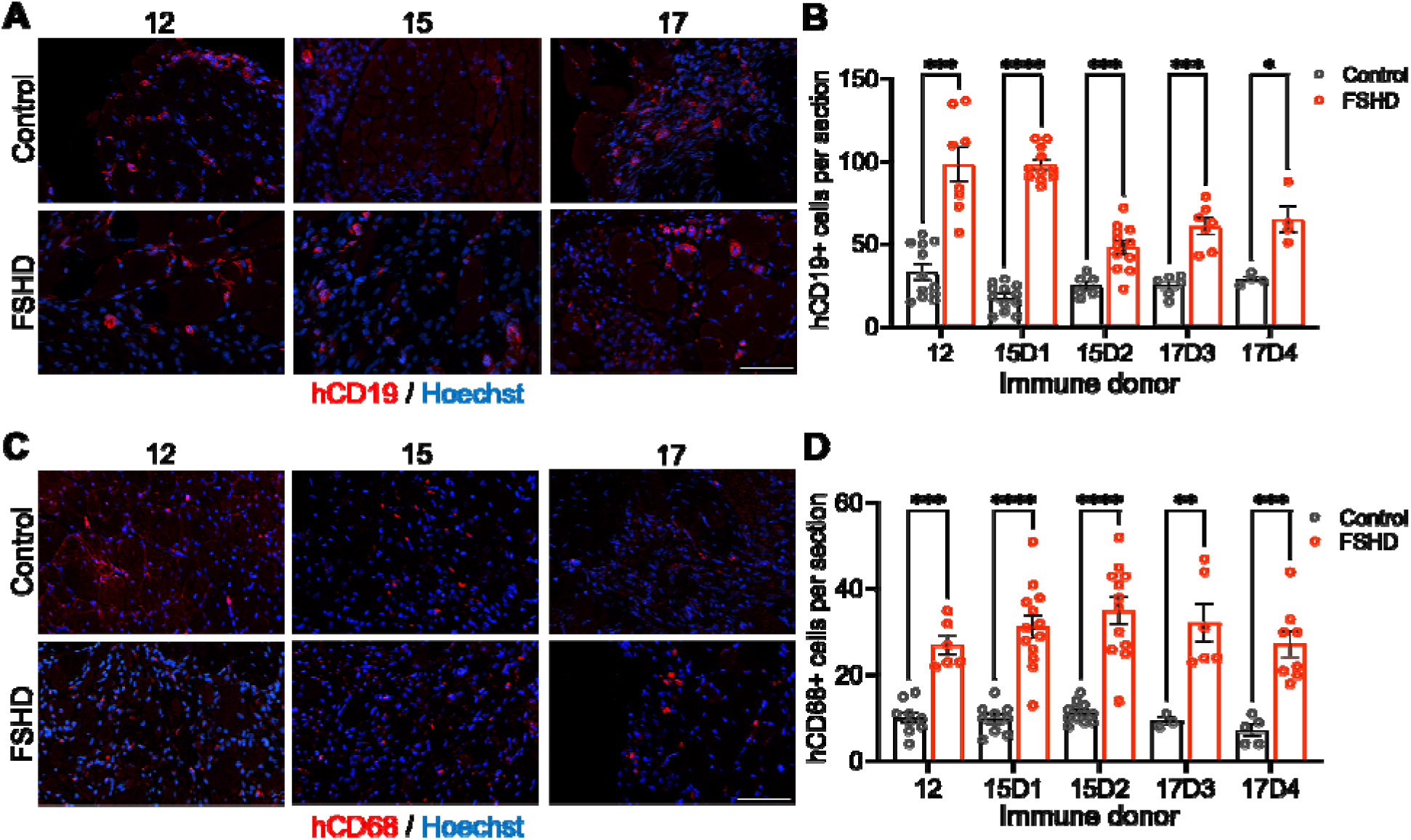
Enhanced accumulation of human B cells and macrophages in FSHD muscle xenografts. Humanized TA muscles were cryosectioned and co-immunostained with human-specific antibodies to CD19 to identify early B cells **(A)** or CD68 to identify macrophages **(C)** and Hoechst to show nuclei for FSHD or control engrafted TAs from mice engrafted with muscle cohorts 12, 15 or 17. Scale bars = 50 μm. **(B)** Quantification of B cell as shown by human CD19 immunostaining. Each dot represents one muscle section with bars showing mean per condition ± SEM. **(D)** Quantification of macrophages as shown by human CD68 immunostaining. Each dot represents one muscle section and is shown as mean per condition ± SEM. *=p<0.05, **=p<0.01, ***=p<0.001, ****=p<0.0001 by Welch’s t-test.

### Human CD19+ B cells and CD68+ macrophages are more abundant in FSHD than control muscle xenografts

To characterize immune cell types in muscle xenografts, FSHD and control TA muscles were sectioned, and serial sections were immunostained with either human CD19 (hCD19) (Figure 4A), a B cell marker, or human CD68 (hCD68) (Figure 4C), a macrophage marker.

Significantly higher numbers of human B cells and macrophages were present in FSHD xenografts compared to control xenografts in all immune donors analyzed (Figure 4B and D). 17D1 and 17D2 TA muscles were processed for IHC and these tissue samples were not suitable for immunofluorescence assays. These data show that FSHD muscle promotes an influx and/or expansion of macrophages and B cells.

### Muscle turnover in FSHD xenografts is immune donor dependent

To investigate whether human innate immune cells promote FSHD muscle turnover, TA muscle sections of FSHD and control xenografts were co-immunostained with human-specific antibodies to lamin A/C to identify human nuclei and spectrin β1 to identify differentiated human muscle fibers in immune donors 12, 15D1, 15D2, 17D3 and 17D4. Xenografts from cohorts 17D1 and 17D2 were processed for IHC and were not assayed for immunofluorescence. Lamin A/C+ nuclei and spectrin β1+ muscle fibers were detected in both FSHD and control TA muscle xenografts (Figure 5A). Notably, FSHD xenografts for cohorts 15D1, 15D2 and 17D4 had significantly fewer spectrin β1+ muscle fibers than control xenografts (Figure 5B), indicating that FSHD muscle is turning over, whereas FSHD and control xenografts from cohorts 12 and 17D3 maintained similar levels of spectrin β1+ muscle fibers, although these FSHD muscle were infiltrated with immune cells (Figure 3 and 4). To confirm that innate immune cell co-engraftment reduced size of FSHD xenografts in engrafted TA muscles, the numbers of spectrin β1+ muscle fibers were compared in immune engrafted mice and mice that failed to develop immune systems in cohorts 12 and 15 (Figure 5C-D).

**Figure 5.**
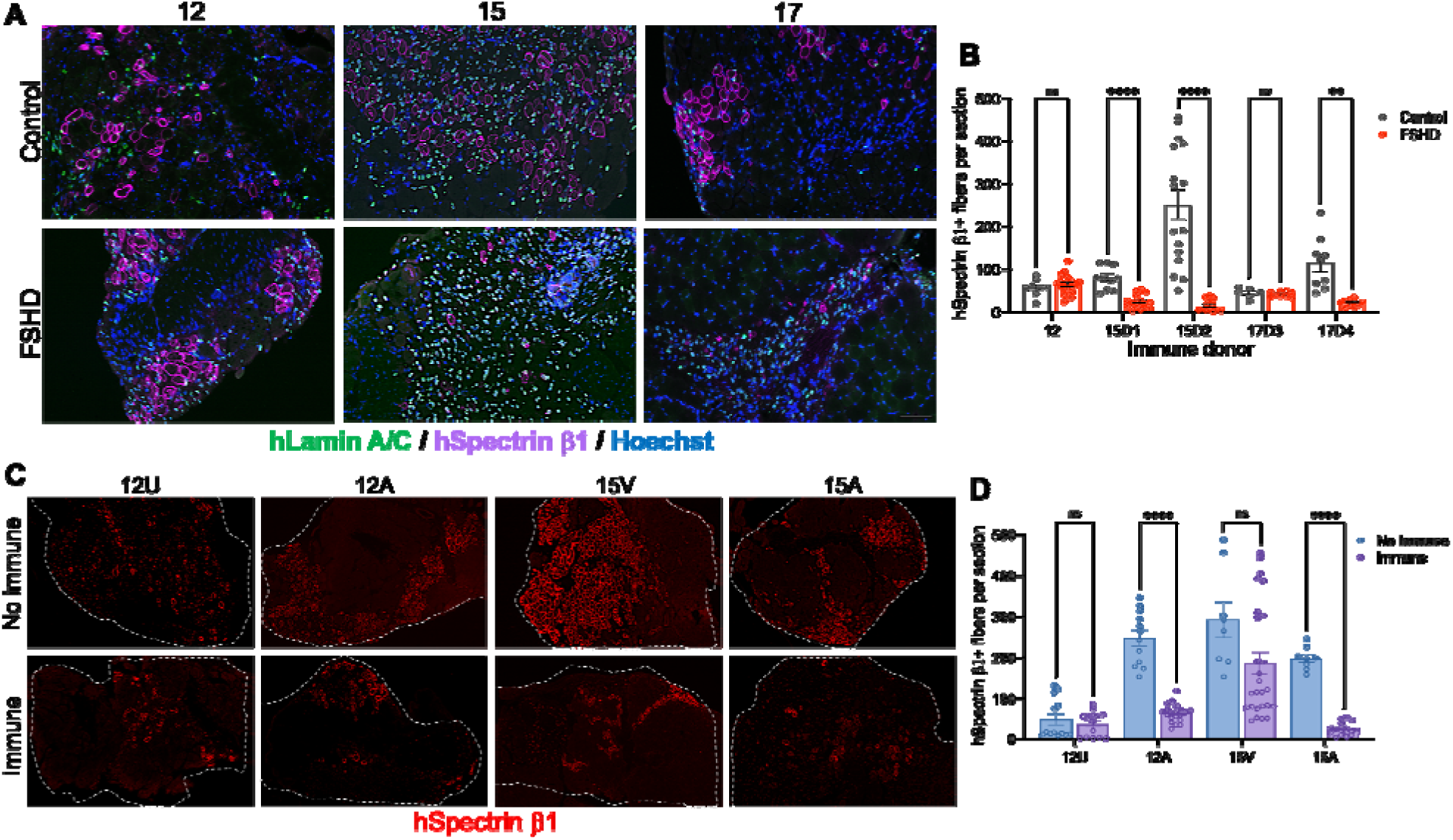
FSHD and unaffected control muscle myoblasts engraft and differentiate in HSC engrafted NSGSGM3-W41 mice. **(A)** Humanized TA muscles were cryosectioned and co-stained with human-specific antibodies to lamin A/C to identify human nuclei, spectrin β1 to identify differentiated muscle fibers and Hoechst to identify all nuclei. Representative images of FSHD and control xenografts from cohorts 12, 15D1, 15D2, 17D3 and 17D4 are shown. Scale bar = 100 μm. **(B)** Quantification of spectrin β1+ fibers for the indicated engraftment conditions. Each dot represents the number of fibers in one muscle section with bars showing mean per condition ± SEM. *=p<0.05, **=p<0.01 by Welch’s t-test. **(C)** Representative images of FSHD and control xenografts from cohorts 12 and 15 with or without immune engraftment stained with spectrin β1 to identify differentiated muscle fibers. Scale bar = 100 μm. **(D)** Quantification of spectrin ®1+ fibers for the indicated engraftment conditions. Each dot represents the number of fibers in one muscle section with bars showing mean per condition ± SEM. ***=p<0.001, ****=p<0.0001 by Welch’s t-test.

Significantly fewer spectrin β1+ muscle fibers were observed in immune engrafted 12A and 15A muscle xenografts, while similar numbers of fibers were observed in 12U and 15V with or without immune engraftment, showing that control muscle does not undergo turnover in response to their co-engrafted immune donor (Figure 5D).

### Inflammatory response to FSHD muscle is immune donor dependent

To analyze the immune and muscle gene expression profiles of FSHD and control xenografts, we designed a custom NanoString RNA expression quantification panel containing human specific probes to immune, muscle and DUX4 target genes. This NanoString panel assayed the expression of 204 inflammation genes, 3 muscle genes (*MYH8*, *MYL2* and *MEF2C*) and 2 DUX4 transcriptional target genes (*LEUTX* and *MBD3L2*). NanoString assays were performed on FSHD and control xenografts from 3 different FSHD families (12, 15 and 17) in NSG- SGM3-W41 mice engrafted with 7 different HSC immune donors, as described above (Figure 2C), and the normalized probe counts for each gene for each mouse can be found in Additional file 2.

To identify differentially expressed genes in FSHD vs control xenografts, NanoString counts for each mouse were log_2_ transformed and averaged within each muscle donor and immune donor combination before calculating the fold change of FSHD vs control expression for all genes within all 7 of the HSC donors. Expression levels of muscle genes were significantly reduced by up to 200-fold (log_2_ fold change -8.37) in 4 of the 7 HSC donor groups (15D1, 15D2, 17D2 and 17D4) (Figure 6A), providing evidence for the differential turnover of FSHD muscle in these cohorts as also observed in immunohistological assays of spectrin β1+ muscle fibers in FSHD xenografts from cohorts 15D1, 15D2 and 17D4 (Figure 5). By contrast, cohorts 12 and 17D3 had increased expression of muscle genes in FSHD vs control, suggestive of a regenerative response of FSHD muscle in these xenografts (Figure 6A). Differentially expressed human immune genes included early complement pathway genes in both the classical and alternative pathways, including *C3*, which is the key mediator of the complement response (45); *C1R*, *C1S*, *C1QA* and *C1QB*, which comprise the C1 complex of the classical complement pathway (46) that initiates complement activation through interactions with pathogens or DAMPs; the *C2* serine proteinase; and *CFB* and *CFD*, which are unique to the alternative pathway and also responsive to DAMPs (Figure 6) (45, 47). Their levels of expression varied based on muscle cohort and HSC immune donor. 17D2 FSHD xenografts had low expression of complement genes and high muscle turnover, but also accumulated macrophages and B cell, suggesting that 17D2 donor immune cells efficiently targeted FSHD muscle for turnover (Figure 4). Notably, *C3* expression trended higher in FSHD xenograft muscles of the three FSHD cohorts responding to all seven immune donors. Expression of all human late complement RNAs including *C5*-*C9,* which encode for components of the Membrane Attack Complex (MAC), were undetectable in both FSHD and control muscles (Additional file 2). Furthermore, NSG-SGM3-W41 mice are deficient in the C5 complement component and therefore cannot elicit a mouse host MAC response. In addition to complement genes, several chemokines including *CXCL1*, *CXCL2*, *CXCL6*, *CXCL9*, *CXCL10* and *CCL13* were expressed higher in FSHD xenografts in several cohorts compared to controls (Additional file 3). The fold changes for all genes in our immune panel can be found in Additional file 3.

**Figure 6:**
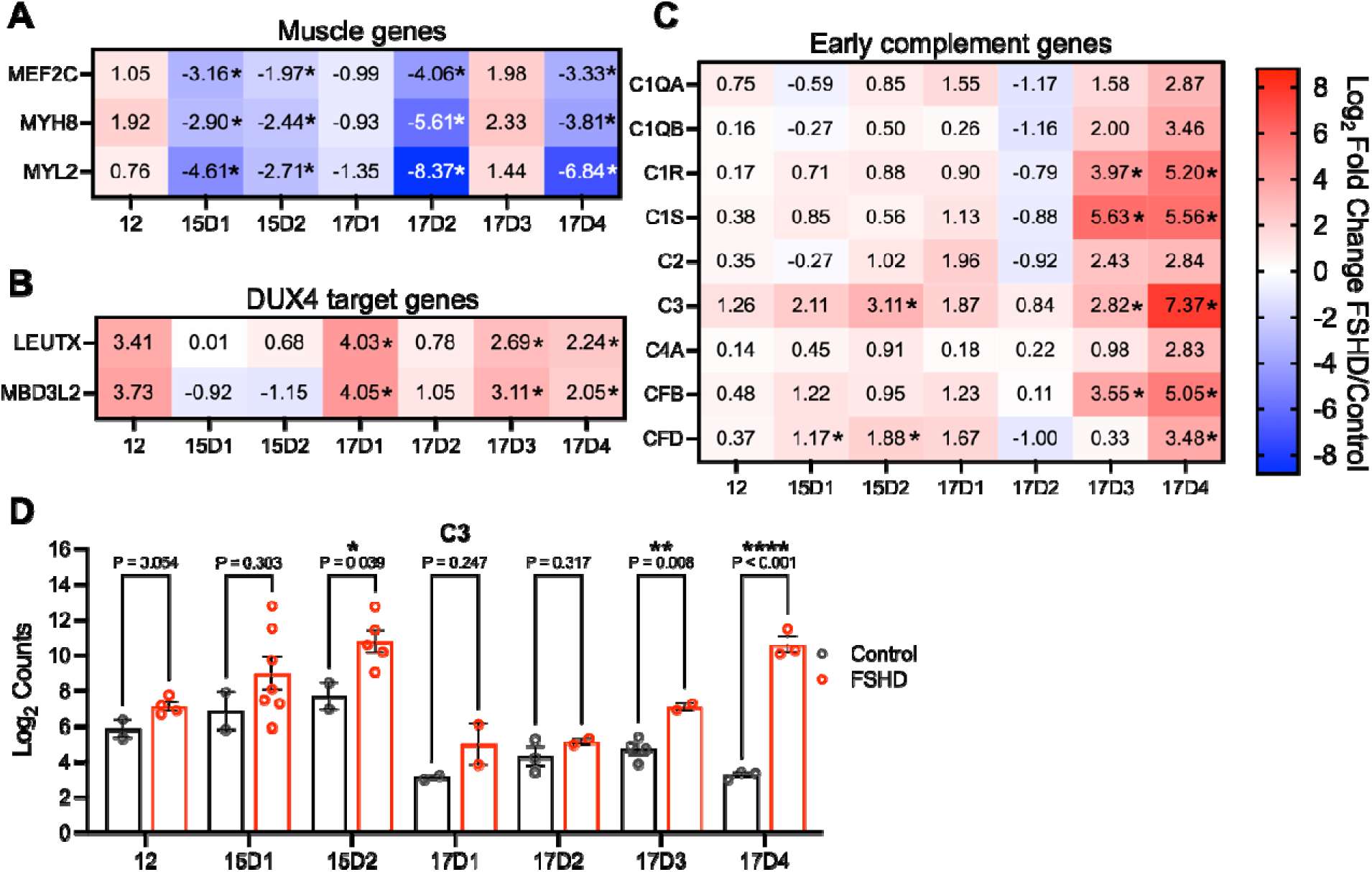
Inflammatory response to FSHD muscle is immune donor dependent. Normalized NanoString counts for muscle genes **(A)**, DUX4 target genes **(B)** and complement genes **(C)** as assayed in RNA isolated from immune/muscle engrafted TA muscles from the indicated cohorts. NanoString counts were log2 transformed and the fold change of FSHD to control gene expression was calculated. Significant gene expression changes are denoted with an asterisk and were calculated using Welch’s t-test. **(D)** NanoString log2 transformed counts showing the expression of C3 RNA from individual TA muscles. Each dot represents expression data from one TA muscle with bars showing mean per condition ± SEM. *=p<0.05, **=p<0.01, ****=p<0.0001 by Welch’s t-test.

### Muscle fibers in FSHD xenografts show increased deposits of C3

Human immune-muscle engrafted TA muscle cryosections were immunostained with human specific antibodies to spectrin β1 to identify human muscle fibers and human specific C3 to investigate the localization of human C3 in relation to humanized muscle regions.

Immunohistology assays of cohorts 12, 15D1, 15D2, 17D3 and 17D4 showed abundant expression and localization of C3 in FSHD muscle fibers compared to control fibers (Figure 7A), supporting the NanoString RNA expression findings. Human C3 was detected only in humanized regions of engrafted mouse TA muscles and localized on the surface and within FSHD muscle fibers but was also concentrated in areas surrounding muscle fibers, which we postulate are enriched in human immune cells. We quantified the abundance of C3 puncta inside spectrin β1+ human muscle fibers and found that in all cohorts analyzed, FSHD engrafted sections had significantly higher percentage of fibers containing greater than 10 C3 puncta (Figure 7B). The high numbers of puncta are highlighted by the white arrows in the high magnification FSHD image (Figure 7A). Mouse C3 was not detected in xenografts by immunohistology, using a mouse specific C3 antibody (data not shown). Taken together, these data demonstrate that FSHD and control xenografts express human C3 and FSHD xenografts have a larger percentage of muscle fibers that are highly decorated with C3.

**Figure 7:**
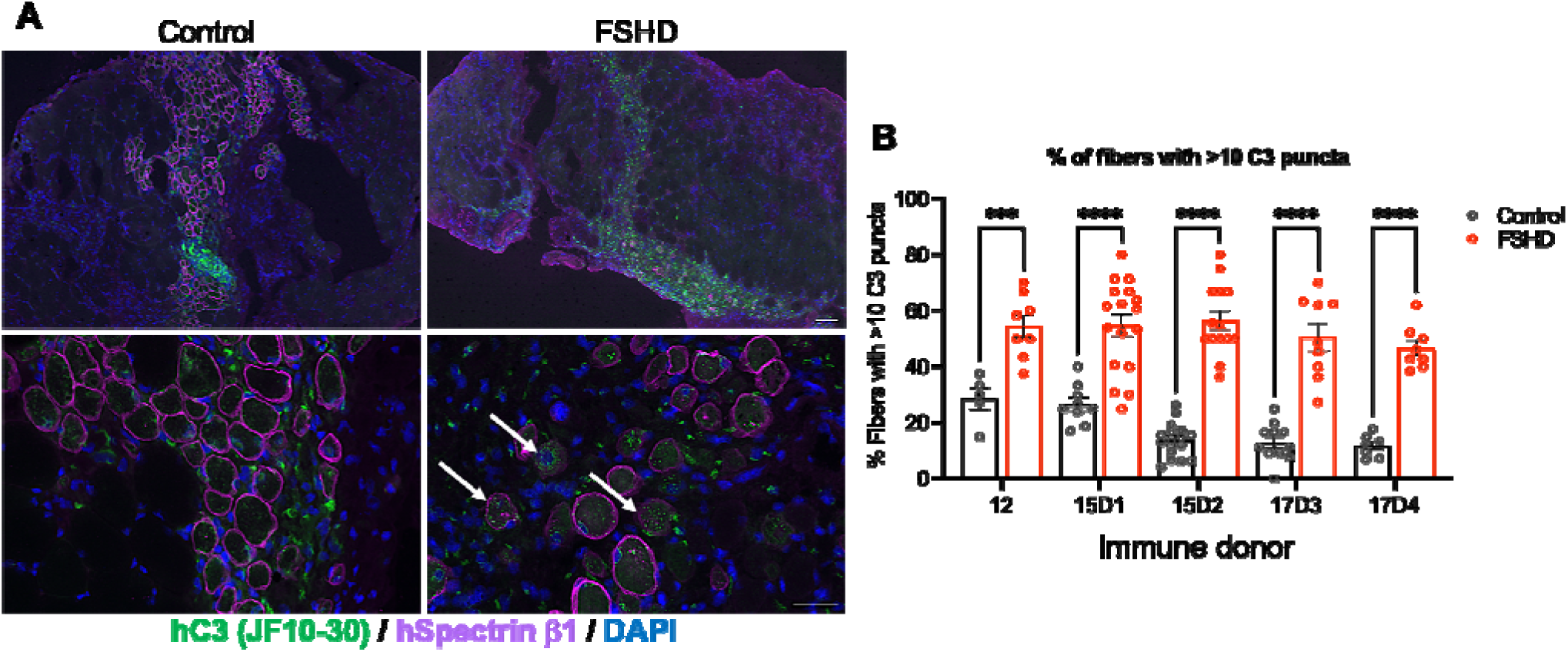
Human C3 localizes to FSHD and control human xenograft muscle fibers. **(A)** Humanized TA muscles were cryosectioned and co-stained with human-specific antibodies to C3, spectrin β1 to identify differentiated muscle fibers and Hoechst to identify nuclei. Representative images of FSHD and control xenografts at low (top) and high magnification (bottom) are shown. Scale bar = (top) 100 μm, (bottom) 50 μm. **(B)** Quantification of C3 puncta in spectrin β1+ fibers for FSHD and control engrafted TA muscle sections from cohorts 12, 15D1, 15D2, 17D3 and 17D4. Each dot represents the percent of fibers with greater than 10 C3 puncta with bars showing mean per condition ± SEM. ***=p<0.001, ****=p<0.0001 by Welch’s t-test.

## Discussion

We have developed a humanized innate immune/muscle mouse model to investigate the role of innate immune responses to FSHD muscle. Our findings establish that FSHD muscle xenografts from all three FSHD families generate innate inflammatory responses to respective immune donors. FSHD xenografts elicited enhanced infiltration of macrophages and B cells compared to control xenografts, showing that FSHD muscle is trophic for and/or promotes the expansion of innate immune cells within FSHD xenografts. FSHD xenografts expressed human early complement RNAs and human C3 RNA and protein as part of an innate immune inflammatory response to FSHD muscle, and the expression of mouse C3 protein could not be detected. FSHD xenografts also underwent what we hypothesize is immune donor-dependent muscle turnover, as shown by the differential muscle turnover responses of 17A FSHD xenografts to four different immune donors (Figure 6). Two of the four 17A immune donors, 17D2 and 17D4 promoted extensive muscle turnover compared to donor 17D1, which promoted lower turnover, and donor 17D3 promoted increased muscle gene expression, perhaps reflecting a regenerative response to muscle damage by the resident muscle myoblasts in xenografts, as observed in FSHD muscle (48). Although muscle turnover in 17D3 was not observed, macrophages and early B cells from both 17D3 and 17D4 HSC donors preferentially accumulated in FSHD muscle xenografts and expressed elevated C3, showing robust inflammatory responses. HSC donor dependent differences in FSHD muscle turnover may reflect quantitative differences in immune donor potency relative to the fixed end point of our assay. Studies are ongoing to establish live animal imaging reporter muscle stem cell lines to establish muscle xenografts. This will enable monitoring of the kinetics of muscle turnover of individual muscle xenografts during stem cell differentiation and maturation in response to different immune donors. Our data demonstrates that individual immune donors produce innate immune cells that have different capacities for immune responses, perhaps modeling aspects of the observed variability in disease progression in FSHD patients and families with multiple affected members (49). Innate immune responses vary in populations, so it is expected that we observe variability in the immune responses from our healthy immune donors in response to FSHD xenografts (50). Future studies of immune donor variability in this model will address these possibilities.

Our innate immune muscle xenograft model establishes a role for innate immunity in FSHD muscle pathology, but the mechanisms by which FSHD muscle is trophic for innate immune cells remain to be investigated. Our working hypothesis is that FSHD muscle attracts macrophages and early B cells through the DUX4-mediated production of DAMPs to stimulate production of complement factor C3 and early complement classical and alternative pathway convertases for processing C3 to C3b. By this mechanism, C3b would bind to and opsonize FSHD muscle for recognition and turnover by macrophage engulfment (45, 47). Current studies are focused on investigations of the functions of DUX4 and C3 in FSHD muscle turnover using DUX4 siRNA therapeutics and early complement pathway-specific immunotherapeutics, with the goal of developing combinatorial therapeutics to treat FSHD disease initiation and progression.

## Conclusions

We have generated a humanized innate immune-FSHD muscle xenograft model using NSG- SGM3-W41 mice to investigate the innate immune response to FSHD muscle. In standardizing our model using muscle biopsy-derived myoblasts from three FSHD patients and paired healthy controls, and HSCs from seven healthy donors, we found that in all cohorts, human B cells and macrophages preferentially infiltrate FSHD muscle. While immune cells infiltrated FSHD xenograft from all cohorts, FSHD muscle turnover was only observed in four of the seven cohorts suggesting that the response is immune donor dependent. Finally, we observed higher expression of complement genes from both the classic and alternative pathways in FSHD engrafted muscles than to control engrafted muscles, suggesting a potential mechanism and novel druggable pathway to ameliorate FSHD muscle pathology.

## List of abbreviations

FSHD: Facioscapulohumeral muscular dystrophy
TA: Tibialis anterior
HSC: hematopoietic stem cells
NMD: non-sense mediated decay
DAMPs: Damage Associated Molecular Patterns
UCB: umbilical cord blood
MAC: Membrane Attack Complex
PBS: phosphate buffered saline.

## Declarations

### Ethics approval and consent to participate

All procedures were performed in accordance with the Institutional Animal Care and Use Committee (IACUC) at the UMass Chan Medical School under protocol number 201900322 and 202000042.

### Availability of data and materials

All data generated or analyzed during this study are included in this published article.

## Acknowledgments

We would like to thank Oliver King and the Emerson and Brehm lab members for critical discussions about data analysis and presentation of results.

## Authors’ contributions

KD and CPE wrote the manuscript. KD, MAB, LDS and CPE edited the manuscript. KD and CPE designed the experiments and analyzed the data. KD, JK and JY performed the experiments. LB, LDS and MAB generated and characterized the NSG-SGM3-W41 mouse strain.

## Additional files

**Additional file 1: Human CD45+ cells in spleen and xenograft muscle of mice engrafted with donor HSC cells. (A)** Representative flow data from HSC and muscle co-engrafted mice including gating strategy to identify human immune cell subtypes are shown. **(B)** Flow cytometry assays from spleen for the percent human CD45+ cells. Animals that failed to develop human immune cell populations are included (no IM). Each dot represents data from one mouse with bars showing mean per condition ± SEM**. (C)** Quantification of immunostaining showing CD45+ cells per section for the indicated engraftment conditions. Each dot represents the number of CD45+ cells in one muscle section with bars showing mean ± SEM per condition.

**Additional file 2: Normalized RNA counts for all genes for each individual mouse TA muscle from all experiments.** Excel file (.xlsx) containing the normalized NanoString counts for each TA analyzed.

**Additional file 3: Expression of inflammatory genes for immune engrafted humanized muscle xenografts for all genes on the custom NanoString panel**. Normalized NanoString counts for inflammation, muscle and DUX4 target genes were assayed for RNA isolated from immune-muscle engrafted TA muscles from the indicated cohorts. Counts were log_2_ transformed and the fold change of FSHD to control gene expression was calculated. Blank spaces are shown for genes that are expressed below the limit of detection for all the animals in the indicated cohort.

**Additional file 4. Expression of immune engrafted humanized muscle compared to non- immune for all genes on the NanoString panel.** Normalized NanoString counts for inflammation, muscle and DUX4 target genes from RNA isolated from immune/muscle engrafted and no immune muscle engrafted TA muscles. Counts from the indicated cohorts were log_2_ transformed and the fold change of immune to no immune gene expression was calculated. Blank spaces are shown for genes that are expressed below the limit of detection for all the animals in the indicated cohort.

## References

1. Jones TI, Chen JC, Rahimov F, Homma S, Arashiro P, Beermann ML, et al. Facioscapulohumeral muscular dystrophy family studies of DUX4 expression: evidence for disease modifiers and a quantitative model of pathogenesis. Hum Mol Genet. 2012;21(20):4419–30.

2. Lemmers RJ, de Kievit P, Sandkuijl L, Padberg GW, van Ommen GJ, Frants RR, et al. Facioscapulohumeral muscular dystrophy is uniquely associated with one of the two variants of the 4q subtelomere. Nat Genet. 2002;32(2):235–6.

3. Lemmers RJ, Wohlgemuth M, van der Gaag KJ, van der Vliet PJ, van Teijlingen CM, de Knijff P, et al. Specific sequence variations within the 4q35 region are associated with facioscapulohumeral muscular dystrophy. Am J Hum Genet. 2007;81(5):884–94.

4. Larsen M, Rost S, El Hajj N, Ferbert A, Deschauer M, Walter MC, et al. Diagnostic approach for FSHD revisited: SMCHD1 mutations cause FSHD2 and act as modifiers of disease severity in FSHD1. Eur J Hum Genet. 2015;23(6):808–16.

5. Lemmers RJ, Tawil R, Petek LM, Balog J, Block GJ, Santen GW, et al. Digenic inheritance of an SMCHD1 mutation and an FSHD-permissive D4Z4 allele causes facioscapulohumeral muscular dystrophy type 2. Nat Genet. 2012;44(12):1370–4.

6. Kowaljow V, Marcowycz A, Ansseau E, Conde CB, Sauvage S, Matteotti C, et al. The DUX4 gene at the FSHD1A locus encodes a pro-apoptotic protein. Neuromuscul Disord. 2007;17(8):611–23.

7. Lemmers RJ, van der Vliet PJ, Klooster R, Sacconi S, Camano P, Dauwerse JG, et al. A unifying genetic model for facioscapulohumeral muscular dystrophy. Science. 2010;329(5999):1650-3.

8. Grow EJ, Weaver BD, Smith CM, Guo J, Stein P, Shadle SC, et al. p53 convergently activates Dux/DUX4 in embryonic stem cells and in facioscapulohumeral muscular dystrophy cell models. Nat Genet. 2021;53(8):1207–20.

9. Hendrickson PG, Dorais JA, Grow EJ, Whiddon JL, Lim JW, Wike CL, et al. Conserved roles of mouse DUX and human DUX4 in activating cleavage-stage genes and MERVL/HERVL retrotransposons. Nat Genet. 2017;49(6):925–34.

10. Geng LN, Yao Z, Snider L, Fong AP, Cech JN, Young JM, et al. DUX4 activates germline genes, retroelements, and immune mediators: implications for facioscapulohumeral dystrophy. Dev Cell. 2012;22(1):38–51.

11. Yao Z, Snider L, Balog J, Lemmers RJ, Van Der Maarel SM, Tawil R, et al. DUX4-induced gene expression is the major molecular signature in FSHD skeletal muscle. Hum Mol Genet. 2014;23(20):5342–52.

12. Wong CJ, Wang LH, Friedman SD, Shaw D, Campbell AE, Budech CB, et al. Longitudinal measures of RNA expression and disease activity in FSHD muscle biopsies. Hum Mol Genet. 2020;29(6):1030–43.

13. Campbell AE, Belleville AE, Resnick R, Shadle SC, and Tapscott SJ. Facioscapulohumeral dystrophy: activating an early embryonic transcriptional program in human skeletal muscle. Hum Mol Genet. 2018;27(R2):R153–R62.

14. Rickard AM, Petek LM, and Miller DG. Endogenous DUX4 expression in FSHD myotubes is sufficient to cause cell death and disrupts RNA splicing and cell migration pathways. Hum Mol Genet. 2015;24(20):5901–14.

15. Jones T, and Jones PL. A cre-inducible DUX4 transgenic mouse model for investigating facioscapulohumeral muscular dystrophy. PLoS One. 2018;13(2):e0192657.

16. Giesige CR, Wallace LM, Heller KN, Eidahl JO, Saad NY, Fowler AM, et al. AAV-mediated follistatin gene therapy improves functional outcomes in the TIC-DUX4 mouse model of FSHD. JCI Insight. 2018;3(22).

17. Jones TI, King OD, Himeda CL, Homma S, Chen JC, Beermann ML, et al. Individual epigenetic status of the pathogenic D4Z4 macrosatellite correlates with disease in facioscapulohumeral muscular dystrophy. Clin Epigenetics. 2015;7:37.

18. Wang LH, Shaw DWW, Faino A, Budech CB, Lewis LM, Statland J, et al. Longitudinal study of MRI and functional outcome measures in facioscapulohumeral muscular dystrophy. BMC Musculoskelet Disord. 2021;22(1):262.

19. Leung DG, Carrino JA, Wagner KR, and Jacobs MA. Whole-body magnetic resonance imaging evaluation of facioscapulohumeral muscular dystrophy. Muscle Nerve. 2015;52(4):512–20.

20. Tasca G, Pescatori M, Monforte M, Mirabella M, Iannaccone E, Frusciante R, et al. Different molecular signatures in magnetic resonance imaging-staged facioscapulohumeral muscular dystrophy muscles. PLoS One. 2012;7(6):e38779.

21. Bosnakovski D, Chan SSK, Recht OO, Hartweck LM, Gustafson CJ, Athman LL, et al. Muscle pathology from stochastic low level DUX4 expression in an FSHD mouse model. Nat Commun. 2017;8(1):550.

22. Mitsuhashi H, Mitsuhashi S, Lynn-Jones T, Kawahara G, and Kunkel LM. Expression of DUX4 in zebrafish development recapitulates facioscapulohumeral muscular dystrophy. Hum Mol Genet. 2013;22(3):568–77.

23. Resnick R, Wong CJ, Hamm DC, Bennett SR, Skene PJ, Hake SB, et al. DUX4-Induced Histone Variants H3.X and H3.Y Mark DUX4 Target Genes for Expression. Cell Rep. 2019;29(7):1812–20 e5.

24. Jae-Hwan Choi Y-EP, Jin-Hong Shin, Chang-Hoon Lee, and Dae-Seong Kim. Extensive inflammatory reaction in facioscapulohumeral muscular dystrophy. Annals of Clinical Neurophysiology. 2017;19(2):141–4.

25. Arahata K, Ishihara T, Fukunaga H, Orimo S, Lee JH, Goto K, et al. Inflammatory response in facioscapulohumeral muscular dystrophy (FSHD): immunocytochemical and genetic analyses. Muscle Nerve Suppl. 1995;2:S56–66.

26. Bosnakovski D, Oyler D, Mitanoska A, Douglas M, Ener ET, Shams AS, et al. Persistent Fibroadipogenic Progenitor Expansion Following Transient DUX4 Expression Provokes a Profibrotic State in a Mouse Model for FSHD. Int J Mol Sci. 2022;23(4).

27. Banerji CRS, Panamarova M, and Zammit PS. DUX4 expressing immortalized FSHD lymphoblastoid cells express genes elevated in FSHD muscle biopsies, correlating with the early stages of inflammation. Hum Mol Genet. 2020;29(14):2285–99.

28. Rahimov F, King OD, Leung DG, Bibat GM, Emerson CP, Jr., Kunkel LM, et al. Transcriptional profiling in facioscapulohumeral muscular dystrophy to identify candidate biomarkers. Proc Natl Acad Sci U S A. 2012;109(40):16234–9.

29. Wang LH, and Tawil R. Facioscapulohumeral Dystrophy. Curr Neurol Neurosci Rep. 2016;16(7):66.

30. Jones TI, Chew GL, Barraza-Flores P, Schreier S, Ramirez M, Wuebbles RD, et al. Transgenic mice expressing tunable levels of DUX4 develop characteristic facioscapulohumeral muscular dystrophy-like pathophysiology ranging in severity. Skelet Muscle. 2020;10(1):8.

31. Feng Q, Snider L, Jagannathan S, Tawil R, van der Maarel SM, Tapscott SJ, et al. A feedback loop between nonsense-mediated decay and the retrogene DUX4 in facioscapulohumeral muscular dystrophy. Elife. 2015;4.

32. Roh JS, and Sohn DH. Damage-Associated Molecular Patterns in Inflammatory Diseases. Immune Netw. 2018;18(4):e27.

33. Wong CJ, Wang L, Holers VM, Frazer-Abel A, van der Maarel SM, Tawil R, et al. Elevated plasma complement components in facioscapulohumeral dystrophy. Hum Mol Genet. 2022;31(11):1821–9.

34. Guo D, Daman K, Chen JJ, Shi MJ, Yan J, Matijasevic Z, et al. iMyoblasts for ex vivo and in vivo investigations of human myogenesis and disease modeling. Elife. 2022;11.

35. Dunkelberger JR, and Song WC. Complement and its role in innate and adaptive immune responses. Cell Res. 2010;20(1):34–50.

36. Bubeck D. The making of a macromolecular machine: assembly of the membrane attack complex. Biochemistry. 2014;53(12):1908–15.

37. Low BE, Kutny PM, and Wiles MV. Simple, Efficient CRISPR-Cas9-Mediated Gene Editing in Mice: Strategies and Methods. Methods Mol Biol. 2016;1438:19–53.

38. Fu Y, Sander JD, Reyon D, Cascio VM, and Joung JK. Improving CRISPR-Cas nuclease specificity using truncated guide RNAs. Nat Biotechnol. 2014;32(3):279–84.

39. Hasgur S, Aryee KE, Shultz LD, Greiner DL, and Brehm MA. Generation of Immunodeficient Mice Bearing Human Immune Systems by the Engraftment of Hematopoietic Stem Cells. Methods Mol Biol. 2016;1438:67–78.

40. Brehm MA, Cuthbert A, Yang C, Miller DM, DiIorio P, Laning J, et al. Parameters for establishing humanized mouse models to study human immunity: analysis of human hematopoietic stem cell engraftment in three immunodeficient strains of mice bearing the IL2rgamma(null) mutation. Clin Immunol. 2010;135(1):84–98.

41. Wunderlich M, Chou FS, Link KA, Mizukawa B, Perry RL, Carroll M, et al. AML xenograft efficiency is significantly improved in NOD/SCID-IL2RG mice constitutively expressing human SCF, GM-CSF and IL-3. Leukemia. 2010;24(10):1785–8.

42. Jangalwe S, Shultz LD, Mathew A, and Brehm MA. Improved B cell development in humanized NOD-scid IL2Rgamma(null) mice transgenically expressing human stem cell factor, granulocyte- macrophage colony-stimulating factor and interleukin-3. Immun Inflamm Dis. 2016;4(4):427–40.

43. McIntosh BE, Brown ME, Duffin BM, Maufort JP, Vereide DT, Slukvin, II, et al. Nonirradiated NOD,B6.SCID Il2rgamma-/- Kit(W41/W41) (NBSGW) mice support multilineage engraftment of human hematopoietic cells. Stem Cell Reports. 2015;4(2):171–80.

44. Rahmig S, Kronstein-Wiedemann R, Fohgrub J, Kronstein N, Nevmerzhitskaya A, Bornhauser M, et al. Improved Human Erythropoiesis and Platelet Formation in Humanized NSGW41 Mice. Stem Cell Reports. 2016;7(4):591–601.

45. Merle NS, Church SE, Fremeaux-Bacchi V, and Roumenina LT. Complement System Part I - Molecular Mechanisms of Activation and Regulation. Front Immunol. 2015;6:262.

46. Arlaud GJ, Gaboriaud C, Thielens NM, Budayova-Spano M, Rossi V, and Fontecilla-Camps JC. Structural biology of the C1 complex of complement unveils the mechanisms of its activation and proteolytic activity. Mol Immunol. 2002;39(7-8):383–94.

47. Merle NS, Noe R, Halbwachs-Mecarelli L, Fremeaux-Bacchi V, and Roumenina LT. Complement System Part II: Role in Immunity. Front Immunol. 2015;6:257.

48. Banerji CRS, Henderson D, Tawil RN, and Zammit PS. Skeletal muscle regeneration in facioscapulohumeral muscular dystrophy is correlated with pathological severity. Hum Mol Genet. 2020;29(16):2746–60.

49. Ruggiero L, Mele F, Manganelli F, Bruzzese D, Ricci G, Vercelli L, et al. Phenotypic Variability Among Patients With D4Z4 Reduced Allele Facioscapulohumeral Muscular Dystrophy. JAMA Netw Open. 2020;3(5):e204040.

50. Randolph HE, Fiege JK, Thielen BK, Mickelson CK, Shiratori M, Barroso-Batista J, et al. Genetic ancestry effects on the response to viral infection are pervasive but cell type specific. Science. 2021;374(6571):1127-33.

