## Supplementary figures and images for "A human immune/muscle xenograft model of FSHD muscle pathology"

### Additional File 1

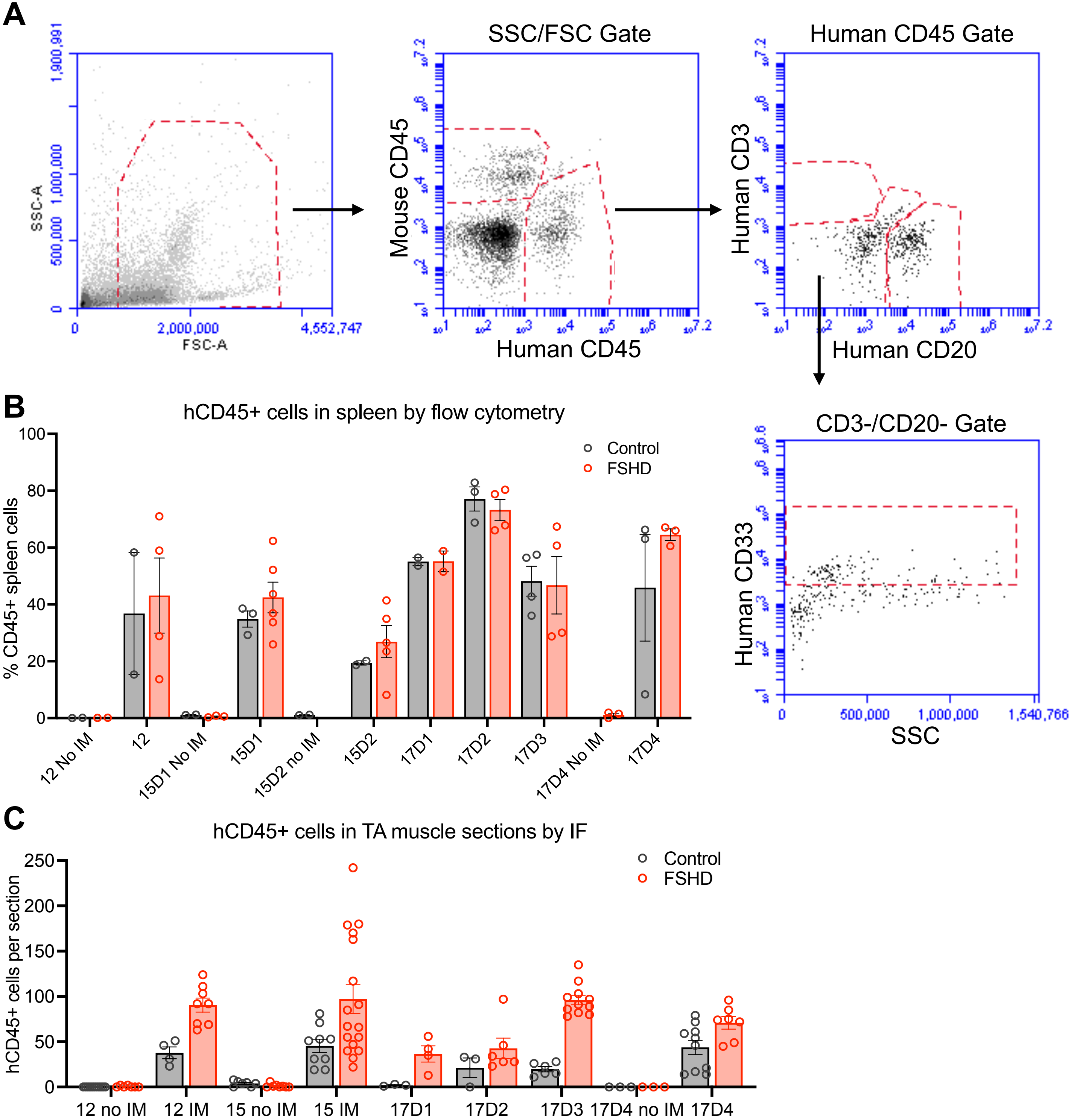
