## Additional File 3 for "A human immune/muscle xenograft model of FSHD muscle pathology"

| Gene | 12 | 15D1 | 15D2 | 17D1 | 17D2 | 17D3 | 17D4 |
|---|---|---|---|---|---|---|---|
| ALOX12 | | | | | | | |
| ALOX15 | | | 0.19 | | | -1.42 | |
| AREG | | | | | | | |
| BCL2L1 | 0.36 | -0.66 | -0.01 | -0.69 | 0.05 | 0.32 | 0.31 |
| BCL6 | 0.26 | -0.18 | 0.47 | | -0.46 | -0.64 | 0.54 |
| BIRC2 | 0.22 | -0.43 | -0.02 | 0.24 | -0.23 | 0.76 | 0.44 |
| C1QA | 0.75 | -0.59 | 0.85 | 1.55 | -1.17 | 1.58 | 2.87 |
| C1QB | 0.16 | -0.27 | 0.50 | 0.26 | -1.16 | 2.00 | 3.46 |
| C1R | 0.17 | 0.71 | 0.88 | 0.90 | -0.79 | 3.97 | 5.20 |
| C1S | 0.38 | 0.85 | 0.56 | 1.13 | -0.88 | 5.63 | 5.56 |
| C2 | 0.35 | -0.27 | 1.02 | 1.96 | -0.92 | 2.43 | 2.84 |
| C3 | 1.26 | 2.11 | 3.11 | 1.87 | 0.84 | 2.82 | 7.37 |
| C3AR1 | | -0.38 | | 1.25 | -0.10 | 0.26 | 0.93 |
| C4A | 0.14 | 0.45 | 0.91 | 0.18 | 0.22 | 0.98 | 2.83 |
| C5 | | | | | | | |
| C6 | | | | | | | |
| C7 | | | | | | | |
| C8A | | | | | | | |
| C8B | | | | | | | |
| C9 | | | | | | | |
| CCL11 | -0.51 | -0.56 | | 0.51 | -0.18 | 0.09 | |
| CCL13 | 0.26 | -0.91 | 1.07 | 1.79 | -0.14 | 1.76 | 3.41 |
| CCL16 | | | | | | | |
| CCL17 | -0.31 | -2.02 | 0.56 | 1.62 | -1.37 | -0.31 | 0.24 |
| CCL19 | -0.19 | -0.80 | 1.44 | | -1.88 | 0.01 | 0.68 |
| CCL2 | 0.71 | -0.67 | 0.04 | 0.58 | -0.64 | 1.39 | 1.71 |
| CCL20 | | | | | | | |
| CCL21 | | | | | | | |
| CCL22 | 0.18 | -1.31 | 0.45 | 0.65 | -1.23 | -0.02 | 0.17 |
| CCL23 | -0.63 | 0.01 | 0.86 | | 0.73 | -0.27 | 0.94 |
| CCL24 | 0.08 | -0.25 | 2.21 | | -3.75 | | 0.85 |
| CCL3 | 0.45 | -0.47 | 1.58 | | 0.09 | | 0.97 |
| CCL4 | 0.14 | -0.78 | 1.57 | 1.02 | 0.46 | -0.54 | 1.02 |
| CCL5 | -0.10 | -0.03 | 1.65 | | -1.30 | 0.34 | 1.24 |
| CCL7 | | | | | | | |
| CCL8 | 0.38 | -0.66 | 1.21 | 1.44 | 0.58 | 0.51 | 2.24 |
| CCR1 | -0.17 | -0.41 | 1.05 | 0.55 | -0.38 | 0.23 | 1.20 |
| CCR2 | -0.22 | -0.33 | 0.31 | | -0.28 | -0.25 | 0.73 |
| CCR3 | -0.19 | | 0.86 | | 0.34 | -0.28 | 0.94 |
| CCR4 | | | | | | | |
| CCR7 | -0.53 | -0.34 | 0.33 | | -0.09 | -0.28 | 0.89 |
| CD163 | 0.45 | -1.16 | 1.01 | 1.40 | -1.17 | 3.45 | 3.64 |
| CD4 | 0.28 | -0.42 | 0.82 | 1.47 | -0.64 | 0.41 | 1.54 |
| CD40 | -0.32 | -0.47 | 0.85 | | -0.71 | -0.60 | 1.58 |
| CD40LG | | | | | | | |
| CD55 | -0.05 | -0.35 | 0.43 | | | -0.42 | |
| CD86 | 0.16 | -0.83 | 0.40 | | -0.65 | -0.30 | 1.42 |
| CEBPB | 0.68 | -0.68 | 0.36 | 0.28 | -1.04 | 0.80 | 0.20 |
| CFB | 0.48 | 1.22 | 0.95 | 1.23 | 0.11 | 3.19 | 5.05 |
| CFD | 0.37 | 1.17 | 1.88 | 1.67 | -1.00 | 1.08 | 3.48 |
| CRP | | | | | | -2.65 | |

| Gene | 12 | 15D1 | 15D2 | 17D1 | 17D2 | 17D3 | 17D4 |
|---|---|---|---|---|---|---|---|
| CSF1 | | | | | | | |
| CSF2 | | | | | | | |
| CSF3 | | 0.10 | | | | | |
| CXCL1 | 0.29 | 1.16 | 0.94 | 1.32 | | 1.77 | 3.61 |
| CXCL10 | -0.07 | -0.02 | 1.08 | 2.19 | -0.53 | -0.43 | 3.19 |
| CXCL2 | 0.13 | 0.09 | 1.80 | | 0.01 | 1.25 | 3.02 |
| CXCL3 | | 0.09 | 1.65 | | 0.02 | 0.31 | 1.37 |
| CXCL5 | | 0.93 | 2.72 | | -1.72 | | |
| CXCL6 | -0.05 | 1.82 | 0.38 | 1.17 | 0.50 | -0.11 | 2.58 |
| CXCL9 | -0.16 | -0.04 | 1.38 | | 0.21 | -0.68 | 2.59 |
| CXCR1 | | | | | | | |
| CXCR2 | -0.28 | -0.03 | 0.32 | | 0.32 | -1.02 | |
| CXCR4 | 0.38 | -1.39 | -0.16 | -0.03 | -1.11 | 0.13 | -0.05 |
| CYSLTR1 | 0.72 | -1.58 | 0.26 | -0.17 | -1.06 | -0.51 | -0.38 |
| CYSLTR2 | | | | | | | |
| DAXX | 0.22 | 0.01 | 0.48 | 0.68 | 0.17 | 0.31 | 0.68 |
| DEFA1 | -1.72 | -3.98 | 0.08 | -0.44 | 1.82 | 1.27 | 0.70 |
| ELK1 | 0.06 | -0.32 | 0.08 | 0.59 | 0.18 | 0.37 | |
| FASLG | | | | | | | |
| FLT1 | 0.04 | -0.10 | 0.55 | | -0.91 | -1.71 | 0.86 |
| FOS | 0.10 | -0.13 | 0.14 | 0.33 | 0.20 | 0.41 | 1.35 |
| FXYD2 | | | | | | | |
| GNB1 | 0.29 | 0.10 | -0.20 | 0.17 | -0.69 | 0.80 | 0.81 |
| GNGT1 | | | | | | | |
| GRB2 | | | | | | | |
| HDAC4 | 0.17 | -0.38 | 0.05 | 0.25 | -0.71 | 1.54 | 0.44 |
| HIF1A | 0.38 | -0.24 | 0.07 | 0.28 | -0.93 | 1.52 | 0.46 |
| HLA-DRA | 0.35 | -1.41 | 0.32 | 1.57 | -1.39 | 0.12 | 2.14 |
| HLA-DRB1 | 0.31 | -1.24 | 0.64 | 1.65 | -1.04 | -0.10 | 2.00 |
| HMGB1 | 0.66 | -0.06 | -0.16 | 0.08 | -0.43 | 0.90 | 0.65 |
| HMGN1 | 0.43 | -0.36 | 0.11 | -0.17 | -0.81 | 0.76 | -0.32 |
| HSH2D | -0.78 | 0.32 | 0.71 | 0.22 | -0.33 | -0.72 | 0.60 |
| IFI44 | 0.17 | | 0.06 | 0.65 | -0.18 | 1.14 | 0.89 |
| IFIT1 | 0.57 | -0.34 | -0.01 | 0.09 | -0.32 | 1.00 | 0.32 |
| IFIT2 | -0.03 | 2.17 | 1.53 | 0.27 | -1.01 | 0.80 | 2.42 |
| IFIT3 | 0.47 | 0.22 | 0.60 | 0.53 | 0.48 | 0.39 | 1.13 |
| IFNA1 | | | | | | | |
| IFNB1 | | | | | | | |
| IFNG | | | | | | | |
| IL10 | | | | | | | |
| IL10RB | 0.37 | 0.18 | 0.05 | 0.56 | -0.43 | 0.98 | 1.35 |
| IL11 | | | | | | | |
| IL12A | | | | | | | |
| IL12B | | | | | | | |
| IL13 | -0.36 | | | | | -1.86 | |
| IL15 | -0.28 | -0.81 | 0.86 | | -0.14 | -0.20 | 1.09 |
| IL17A | | | | | | | |
| IL18 | 0.24 | -0.70 | 0.18 | | -0.93 | -0.45 | 0.63 |
| IL18RAP | -1.48 | -0.40 | 1.00 | | 0.10 | -1.24 | |
| IL1A | | | | | | | |
| IL1B | | | 1.47 | | 0.15 | | |

| Gene | 12 | 15D1 | 15D2 | 17D1 | 17D2 | 17D3 | 17D4 |
|---|---|---|---|---|---|---|---|
| IL1R1 | 0.11 | 1.19 | 0.89 | 0.71 | -0.47 | 1.32 | 2.52 |
| IL1RN | 0.40 | -0.50 | 1.37 | | -1.06 | -1.04 | 1.66 |
| IL2 | -0.41 | | | | | | |
| IL21 | | | | | | | |
| IL22 | | | | | | | |
| IL22RA2 | | | | | | | |
| IL23A | | | | | | | |
| IL3 | -0.14 | 1.05 | -0.12 | 0.66 | 1.69 | -4.04 | 0.09 |
| IL4 | -1.05 | | | | | -1.58 | |
| IL5 | -1.21 | | | | | | |
| IL6 | | | | | | | |
| IL6R | -0.48 | 0.07 | 0.74 | | -0.41 | -0.59 | 1.06 |
| IL7 | | | 1.20 | | | | |
| IL8 | | 0.02 | 1.69 | | -1.37 | 0.14 | 2.24 |
| IL9 | -1.71 | | | | | | |
| IRF1 | -0.10 | -0.02 | 1.02 | | 0.28 | -0.38 | 1.23 |
| IRF3 | -0.15 | -0.13 | 0.27 | | 0.87 | 0.42 | 1.72 |
| IRF5 | -0.40 | -0.22 | 0.26 | 1.05 | -0.53 | -0.31 | 1.26 |
| IRF7 | -0.10 | -0.35 | 0.18 | 0.36 | -0.16 | 0.22 | 0.47 |
| ITGB2 | -0.10 | -0.29 | 1.05 | 1.58 | -1.05 | -0.01 | 1.55 |
| JUN | 0.40 | -0.36 | -0.12 | 0.07 | -0.57 | 1.64 | |
| KNG1 | | | | | | | |
| LEUTX | 3.41 | 0.23 | 0.68 | 4.03 | 0.78 | 2.69 | 2.24 |
| LIMK1 | 0.68 | -0.09 | -0.11 | | -1.19 | 1.02 | 0.42 |
| LTA | | | | | | | |
| LTB | -0.52 | -0.32 | 0.45 | 1.51 | 0.80 | -1.06 | 1.10 |
| LTB4R | -1.04 | -0.23 | 0.52 | -0.05 | 0.43 | -1.56 | |
| LTB4R2 | -0.96 | 0.38 | 0.23 | | 0.18 | 0.22 | |
| MAFF | -0.69 | 0.30 | 0.44 | | | -0.95 | |
| MAFG | 0.20 | -0.59 | 0.29 | 0.63 | -0.47 | 0.40 | 0.24 |
| MAFK | -0.22 | -0.41 | 0.31 | 0.54 | -0.68 | 0.79 | 0.73 |
| MAP2K4 | 0.46 | -0.62 | -0.26 | 0.13 | 0.11 | 0.50 | -0.40 |
| MAP3K9 | | | | | | | |
| MASP1 | | | | | | | |
| MBD3L2 | 3.73 | -0.83 | -1.15 | 4.05 | 1.05 | 3.11 | 2.05 |
| MBL2 | | | | | | | |
| MEF2B | | | | | | | |
| MEF2C | 1.05 | -2.94 | -1.97 | -0.99 | -4.06 | 1.98 | -3.33 |
| MEF2D | -0.20 | -0.84 | -0.21 | -0.14 | -1.18 | 0.87 | 0.54 |
| MKNK1 | 0.09 | -0.40 | 0.65 | 0.88 | -0.88 | 1.10 | 1.05 |
| MMP3 | | | | | | | |
| MMP9 | 0.12 | 0.08 | 2.59 | 0.10 | -0.97 | 2.00 | 2.75 |
| MRC1 | 0.50 | -1.02 | 0.71 | 1.71 | -1.68 | 1.68 | 2.91 |
| MX1 | -0.38 | 0.21 | 0.29 | 0.84 | -0.18 | 0.34 | 0.96 |
| MX2 | -0.14 | 0.61 | 0.70 | 0.54 | 0.09 | 0.64 | 1.52 |
| MYD88 | 0.13 | 0.20 | 0.21 | 1.02 | -0.97 | 0.14 | 1.12 |
| MYH8 | 1.92 | -2.65 | -2.44 | -0.93 | -5.61 | 2.33 | -3.81 |
| MYL2 | 0.76 | -4.25 | -2.71 | -1.35 | -8.37 | 1.44 | -6.84 |
| NFE2L2 | 0.42 | -0.26 | 0.28 | 0.42 | -0.67 | 1.34 | 0.64 |
| NFKB1 | 0.36 | -0.03 | 0.07 | 0.06 | -0.56 | 0.21 | 0.59 |
| NLRP3 | | -0.63 | 0.43 | | -0.58 | 0.25 | 1.17 |

| Gene | 12 | 15D1 | 15D2 | 17D1 | 17D2 | 17D3 | 17D4 |
|---|---|---|---|---|---|---|---|
| NOD1 | | | 0.47 | 0.38 | 0.09 | -0.08 | 1.63 |
| NOD2 | 0.50 | -0.60 | 0.33 | 1.09 | -1.42 | 0.01 | 0.73 |
| NOS2 | | | | | | | |
| NOX1 | | | | | | | |
| OAS2 | 0.05 | 0.30 | 0.14 | 0.71 | -0.41 | 0.81 | 0.84 |
| OASL | | 0.52 | 0.19 | | 0.17 | -0.49 | |
| OXER1 | | | 1.06 | | | | |
| PDGFA | | -1.02 | -0.44 | | | -0.10 | |
| PPP1R12B | 0.70 | -2.57 | -0.25 | -0.20 | -2.30 | 1.92 | -1.89 |
| PRKCA | 0.24 | -0.10 | 0.51 | | -0.31 | -0.04 | 0.98 |
| PTGDR2 | -0.97 | 0.08 | 1.03 | | 1.28 | -1.52 | 1.14 |
| PTGER2 | -0.09 | 0.17 | 0.73 | 0.39 | 0.21 | -0.37 | 1.26 |
| PTGER3 | -0.70 | -0.41 | 0.61 | 0.19 | 0.91 | -1.26 | 0.47 |
| PTGIR | | | | | | | |
| PTGS1 | 0.62 | -0.24 | 0.80 | 1.10 | 0.53 | -0.89 | 1.16 |
| PTGS2 | | 1.23 | 0.87 | | | -0.17 | 2.99 |
| RAC1 | 0.05 | -0.27 | -0.29 | -0.08 | -0.59 | 1.09 | 0.51 |
| RAF1 | 0.53 | -1.14 | 0.04 | -0.03 | -1.05 | 1.04 | -0.31 |
| RAPGEF2 | 0.01 | 0.09 | -0.09 | 0.57 | 0.45 | -0.60 | 0.41 |
| RELA | -0.12 | -0.10 | 0.06 | 0.13 | -0.42 | 0.75 | 0.97 |
| RELB | 0.19 | -0.32 | | 0.15 | 0.23 | 0.05 | 0.50 |
| RHOA | 0.40 | -0.21 | -0.07 | 0.27 | -0.84 | 0.95 | 0.07 |
| RIPK1 | 0.08 | 0.26 | -0.13 | 0.49 | -0.28 | 0.55 | 1.11 |
| RIPK2 | 0.22 | 0.08 | 0.62 | | | | 0.83 |
| SHC1 | 0.53 | 0.24 | | 0.33 | -0.62 | 1.90 | 0.69 |
| SMAD7 | 0.05 | 0.27 | 0.07 | 0.08 | -0.16 | 0.27 | 0.30 |
| STAT1 | 0.22 | -0.42 | 0.05 | 0.49 | -0.51 | 1.49 | 1.26 |
| STAT2 | -0.12 | 0.21 | 0.28 | 0.60 | 0.05 | 0.92 | 1.63 |
| STAT3 | 0.13 | -0.24 | 0.17 | 0.44 | -0.62 | 0.97 | 1.07 |
| TBXA2R | | | | | | | |
| TCF4 | 0.15 | -0.13 | -0.19 | 0.30 | -1.02 | 2.52 | 1.26 |
| TGFB1 | -0.36 | 0.15 | 0.53 | 0.88 | | 0.17 | 1.02 |
| TGFB2 | 0.75 | -1.39 | -0.40 | -0.41 | -2.02 | 1.68 | -1.15 |
| TGFB3 | 0.70 | -0.71 | -0.41 | -0.22 | -4.32 | 3.27 | -0.47 |
| TGFBR1 | -0.02 | 0.02 | 0.54 | 0.79 | -0.20 | -0.01 | 0.85 |
| TLR1 | 0.05 | -0.45 | 0.33 | 1.19 | -1.00 | 0.33 | 1.76 |
| TLR2 | 0.55 | -0.36 | 0.37 | 1.11 | -1.27 | 0.35 | 2.52 |
| TLR3 | 0.26 | 0.07 | -0.03 | | | 0.97 | 1.14 |
| TLR4 | 0.36 | 0.01 | 0.52 | 0.48 | -0.85 | 0.97 | 2.17 |
| TLR5 | | -0.82 | | | | | |
| TLR6 | -0.15 | 0.29 | 0.54 | | -0.91 | -0.33 | 1.29 |
| TLR7 | | -1.16 | | | | | |
| TLR8 | 0.19 | -0.75 | 0.75 | | -1.35 | 0.21 | 1.73 |
| TLR9 | | | | | | | |
| TNF | -0.15 | -0.50 | 1.08 | 1.34 | -0.06 | -1.00 | 1.21 |
| TNFAIP3 | 0.52 | -1.03 | -0.02 | 0.28 | -0.12 | 0.86 | 0.26 |
| TNFSF14 | | | | | | | |
| TOLLIP | 0.20 | -0.30 | 0.08 | 0.36 | -0.37 | 0.73 | 0.81 |
| TRADD | 0.55 | 0.24 | 0.28 | | -0.62 | 0.35 | 0.66 |
| TRAF2 | | 0.18 | 0.53 | | | 0.17 | |
| TREM2 | | | | | | | |
| TSLP | | | | | | | |
| TWIST2 | | | | | | | |
| TYROBP | 0.50 | -1.14 | 0.37 | 1.29 | -1.01 | 0.15 | 1.40 |

$Log_2$ Fold Change FSHD/Control (scale: 8, 6, 4, 2, 0, -2, -4, -6, -8)
