## Additional File 4 for "A human immune/muscle xenograft model of FSHD muscle pathology"

| Gene | 12U | 12A | 15VD1 | 15AD1 | 17AD4 |
|---|---|---|---|---|---|
| ALOX12 | | | | | |
| ALOX15 | | | | | |
| AREG | | | | | |
| BCL2L1 | -0.03 | 0.12 | -0.24 | 0.47 | 0.93 |
| BCL6 | 0.96 | 0.70 | -0.40 | 0.80 | 1.81 |
| BIRC2 | -0.44 | -0.17 | -0.12 | -0.19 | 0.47 |
| C1QA | 4.68 | 4.50 | 4.01 | 7.13 | 7.80 |
| C1QB | 6.03 | 4.83 | 4.57 | 4.92 | 7.22 |
| C1R | 0.96 | 0.59 | 1.09 | 1.84 | 2.34 |
| C1S | 0.05 | 0.35 | 0.95 | 1.18 | 1.88 |
| C2 | 4.10 | 3.84 | 3.50 | 4.03 | 6.45 |
| C3 | 2.93 | 3.74 | 2.93 | 4.19 | 5.70 |
| C3AR1 | 5.07 | 3.95 | 2.90 | 3.62 | 6.74 |
| C4A | 2.67 | 0.93 | 1.06 | 2.10 | 3.00 |
| C5 | | | | | |
| C6 | | | | | |
| C7 | | | | | |
| C8A | | | | | |
| C8B | | | | | |
| C9 | | | | | |
| CCL11 | 1.14 | -0.28 | 0.51 | 0.09 | |
| CCL13 | 7.03 | 5.12 | 3.08 | 4.77 | 8.69 |
| CCL16 | | | | | |
| CCL17 | 7.00 | 4.80 | 5.56 | 4.52 | 5.64 |
| CCL19 | 2.68 | 2.27 | 2.86 | 2.33 | 3.22 |
| CCL2 | 0.62 | 1.26 | 0.65 | 1.60 | 3.68 |
| CCL20 | | | | | |
| CCL21 | | | | | |
| CCL22 | 4.89 | 3.20 | 3.64 | 2.61 | 3.57 |
| CCL23 | 4.87 | 3.40 | 2.35 | 3.42 | 5.48 |
| CCL24 | 3.73 | 2.43 | 3.03 | 2.75 | 4.01 |
| CCL3 | 2.83 | 1.82 | 2.08 | 2.84 | 4.13 |
| CCL4 | 4.16 | 2.63 | 2.36 | 2.14 | 4.27 |
| CCL5 | 2.01 | 2.29 | 2.60 | 2.63 | 3.24 |
| CCL7 | | | | | |
| CCL8 | 2.73 | 2.06 | 2.48 | 3.48 | 4.21 |
| CCR1 | 3.15 | 1.99 | 2.90 | 2.13 | 4.34 |
| CCR2 | 3.51 | 2.24 | 2.18 | 1.58 | 3.63 |
| CCR3 | 2.95 | 1.58 | | | 3.30 |
| CCR4 | | | | | |
| CCR7 | 3.06 | 2.68 | 0.53 | 1.81 | 3.41 |
| CD163 | 4.69 | 4.70 | 3.89 | 3.90 | 8.00 |
| CD4 | 4.15 | 3.84 | 2.60 | 2.76 | 5.05 |
| CD40 | 2.62 | 2.16 | 1.84 | 0.98 | 3.01 |
| CD40LG | | | | | |
| CD55 | 0.15 | -0.42 | -0.44 | -0.31 | |
| CD86 | 4.54 | 4.18 | 3.80 | 3.50 | 5.04 |
| CEBPB | 0.80 | 1.17 | 0.01 | 0.45 | 1.66 |
| CFB | -0.06 | 0.80 | 1.51 | 1.82 | 2.85 |
| CFD | 1.04 | 0.16 | 2.27 | 2.07 | 1.75 |

| Gene | 12U | 12A | 15VD1 | 15AD1 | 17AD4 |
|---|---|---|---|---|---|
| CSF1 | | | | | |
| CSF2 | | | | | |
| CSF3 | | | | | |
| CXCL1 | -0.74 | -0.02 | -0.32 | 1.60 | 3.13 |
| CXCL10 | 2.73 | 0.91 | 5.16 | 3.23 | 4.77 |
| CXCL2 | 1.80 | 1.80 | 2.43 | 1.53 | 3.25 |
| CXCL3 | | | 2.35 | 2.50 | 2.69 |
| CXCL5 | | | 1.58 | 0.86 | |
| CXCL6 | 0.26 | -0.07 | -1.15 | 0.98 | 1.61 |
| CXCL9 | 2.32 | 0.09 | 2.45 | 0.62 | 3.24 |
| CXCR1 | | | | | |
| CXCR2 | 2.52 | 0.92 | 0.60 | 0.51 | |
| CXCR4 | 3.73 | 1.87 | 0.21 | 3.15 | 2.40 |
| CYSLTR1 | 1.25 | 0.87 | 0.46 | 1.00 | 2.05 |
| CYSLTR2 | | | | | |
| DAXX | -0.62 | 0.12 | 0.11 | 0.33 | 0.82 |
| DEFA1 | 4.38 | 2.06 | 5.83 | 1.61 | 1.65 |
| ELK1 | -0.15 | -0.17 | -0.17 | -0.15 | |
| FASLG | | | | | |
| FLT1 | 3.35 | 1.83 | 2.30 | 1.96 | 3.57 |
| FOS | -0.32 | 0.61 | 0.31 | -0.50 | 0.63 |
| FXYD2 | | | | | |
| GNB1 | -0.81 | -0.20 | -0.16 | -0.05 | 0.21 |
| GNGT1 | | | | | |
| GRB2 | | | | | |
| HDAC4 | -0.07 | -0.26 | -0.65 | -0.28 | -0.23 |
| HIF1A | -1.15 | -0.27 | -0.21 | -0.01 | 0.25 |
| HLA-DRA | 8.40 | 7.05 | 4.16 | 6.13 | 9.50 |
| HLA-DRB1 | 8.28 | 7.03 | 3.43 | 5.59 | 9.23 |
| HMGB1 | -1.34 | -0.22 | -0.06 | -0.12 | 0.01 |
| HMGN1 | -0.96 | -0.28 | -0.23 | -0.01 | 0.10 |
| HSH2D | 3.46 | 2.04 | 1.91 | 1.17 | 2.62 |
| IFI44 | 1.53 | 1.67 | 1.49 | 2.47 | 2.65 |
| IFIT1 | 0.89 | 1.46 | 1.87 | 2.56 | 1.94 |
| IFIT2 | 1.27 | 0.92 | 1.09 | 2.26 | 1.76 |
| IFIT3 | 1.00 | 1.37 | 2.37 | 2.01 | 2.58 |
| IFNA1 | | | | | |
| IFNB1 | | | | | |
| IFNG | | | | | |
| IL10 | | | | | |
| IL10RB | -0.44 | 0.40 | 0.32 | 0.42 | 0.91 |
| IL11 | | | | | |
| IL12A | | | | | |
| IL12B | | | | | |
| IL13 | 1.63 | 0.74 | | | |
| IL15 | 1.46 | 1.80 | 0.84 | 0.81 | 1.90 |
| IL17A | | | | | |
| IL18 | 1.83 | 0.21 | -0.07 | 0.89 | 3.63 |
| IL18RAP | 3.60 | 0.98 | 2.68 | 2.71 | |

| Gene | 12U | 12A | 15VD1 | 15AD1 | 17AD4 |
|---|---|---|---|---|---|
| IL1R1 | -0.71 | -0.38 | -0.11 | -0.02 | 0.36 |
| IL1RN | 2.35 | 2.11 | 2.39 | 4.17 | 3.82 |
| IL2 | 1.17 | 0.22 | | | |
| IL21 | | | | | |
| IL22 | | | | | |
| IL22RA2 | | | | | |
| IL23A | | | | | |
| IL3 | -0.20 | 0.12 | -1.00 | -0.65 | -1.26 |
| IL4 | 2.40 | 0.45 | | | |
| IL5 | 1.60 | 1.61 | | | |
| IL6 | | | | | |
| IL6R | 2.36 | 1.13 | 0.74 | -0.14 | 2.03 |
| IL7 | | | | | |
| IL8 | | | 1.70 | 1.88 | 4.10 |
| IL9 | 1.34 | | | | |
| IRF1 | 1.23 | 0.68 | 0.43 | 0.60 | 2.66 |
| IRF3 | 0.09 | -0.12 | 0.12 | -0.16 | 0.76 |
| IRF5 | 3.57 | 1.91 | 2.86 | 2.26 | 4.46 |
| IRF7 | 1.53 | 1.02 | 0.37 | 1.91 | 2.58 |
| ITGB2 | 5.51 | 4.96 | 3.25 | 3.81 | 6.62 |
| JUN | -1.25 | -0.15 | -0.68 | -0.40 | 0.06 |
| KNG1 | | | | | |
| LEUTX | | -0.56 | | -1.49 | -1.40 |
| LIMK1 | -1.35 | -0.26 | -0.16 | -0.19 | -0.14 |
| LTA | | | | | |
| LTB | 4.85 | 3.53 | 3.42 | 1.94 | 4.38 |
| LTB4R | 2.95 | 0.52 | 0.43 | 1.11 | |
| LTB4R2 | 1.59 | -0.47 | 0.08 | 0.46 | |
| MAFF | 2.16 | 1.04 | -0.17 | 0.55 | |
| MAFG | -0.19 | -0.20 | -0.43 | 0.10 | 0.59 |
| MAFK | 0.22 | -0.17 | -0.57 | -0.11 | 0.28 |
| MAP2K4 | -0.79 | -0.41 | -0.25 | -0.04 | -0.23 |
| MAP3K9 | | | | | |
| MASP1 | | | | | |
| MBD3L2 | | -0.68 | 1.45 | -0.63 | -0.76 |
| MBL2 | | | | | |
| MEF2B | | | | | |
| MEF2C | -0.28 | -0.81 | -0.48 | -1.53 | -0.72 |
| MEF2D | -0.04 | -0.44 | -0.52 | -0.65 | 0.59 |
| MKNK1 | 0.29 | 0.44 | 0.66 | 1.15 | 1.44 |
| MMP3 | | | | | |
| MMP9 | 5.06 | 3.45 | 2.31 | 3.95 | 4.73 |
| MRC1 | 6.11 | 4.65 | 3.24 | 4.86 | 8.02 |
| MX1 | 3.41 | 3.83 | 3.10 | 4.27 | 4.07 |
| MX2 | 2.74 | 2.43 | 2.69 | 2.62 | 3.57 |
| MYD88 | 1.17 | 1.66 | 1.30 | 1.65 | 2.01 |
| MYH8 | -1.41 | -1.26 | -0.11 | -1.75 | -1.80 |
| MYL2 | 1.35 | -0.73 | -0.75 | -2.51 | -1.69 |
| NFE2L2 | 1.35 | -0.10 | -0.27 | -0.01 | 0.49 |
| NFKB1 | -0.53 | 0.28 | -0.22 | 0.09 | 0.83 |
| NLRP3 | | | 1.83 | 2.42 | 3.11 |

| Gene | 12U | 12A | 15VD1 | 15AD1 | 17AD4 |
|---|---|---|---|---|---|
| NOD1 | 0.75 | 0.47 | 0.25 | 0.76 | 1.26 |
| NOD2 | 4.37 | 2.36 | 1.71 | 2.02 | 3.87 |
| NOS2 | | | | | |
| NOX1 | | | | | |
| OAS2 | 2.63 | 2.53 | 1.88 | 3.87 | 3.61 |
| OASL | | | 1.14 | 1.42 | |
| OXER1 | | | | | |
| PDGFA | | | -0.50 | -0.59 | |
| PPP1R12B | -0.60 | -0.63 | -0.41 | -1.07 | -0.36 |
| PRKCA | 0.35 | 0.86 | 0.11 | 0.06 | 1.27 |
| PTGDR2 | 2.41 | 0.92 | 1.74 | 1.67 | 0.89 |
| PTGER2 | 1.46 | 1.53 | 0.35 | 0.96 | 1.59 |
| PTGER3 | 3.86 | 2.01 | 1.28 | 1.59 | 4.07 |
| PTGIR | | | | | |
| PTGS1 | 1.80 | 1.48 | 1.77 | 1.46 | 3.52 |
| PTGS2 | | | 0.69 | 0.75 | 1.40 |
| RAC1 | -0.66 | -0.25 | -0.19 | -0.15 | 0.10 |
| RAF1 | -0.57 | -0.11 | -0.31 | -0.39 | 0.89 |
| RAPGEF2 | 0.24 | 0.46 | -0.46 | 0.33 | 1.43 |
| RELA | -0.38 | -0.24 | -0.11 | 0.41 | 0.62 |
| RELB | -0.25 | -0.02 | 0.13 | 0.31 | 1.43 |
| RHOA | -1.01 | -0.34 | -0.18 | -0.05 | 0.05 |
| RIPK1 | -0.52 | 0.14 | 0.15 | 0.21 | 0.06 |
| RIPK2 | -0.25 | 0.34 | 0.29 | 0.85 | 1.11 |
| SHC1 | -1.22 | -0.49 | -0.48 | -0.13 | -0.38 |
| SMAD7 | 0.25 | 0.30 | -0.13 | -0.12 | 0.79 |
| STAT1 | 0.84 | 1.28 | 1.74 | 2.19 | 3.16 |
| STAT2 | 0.32 | 0.12 | 0.40 | 0.85 | 1.19 |
| STAT3 | -0.17 | 0.21 | 0.03 | 0.43 | 0.85 |
| TBXA2R | | | | | |
| TCF4 | -1.13 | -0.68 | -0.33 | -0.61 | -0.55 |
| TGFB1 | 0.89 | 0.71 | 0.02 | 0.59 | 1.81 |
| TGFB2 | -2.05 | -0.62 | -0.46 | -0.98 | -0.93 |
| TGFB3 | -1.94 | -0.88 | -0.79 | -0.95 | -1.45 |
| TGFBR1 | 0.51 | 0.51 | 0.34 | 0.58 | 1.29 |
| TLR1 | 1.50 | 1.53 | 2.00 | 1.95 | 2.72 |
| TLR2 | 3.68 | 2.83 | 2.39 | 2.21 | 4.09 |
| TLR3 | 0.80 | 0.69 | 1.04 | 1.25 | 2.16 |
| TLR4 | 0.24 | 1.21 | 0.52 | 0.99 | 1.64 |
| TLR5 | | | 1.24 | 0.85 | |
| TLR6 | 0.96 | 1.23 | 0.77 | 0.72 | 2.07 |
| TLR7 | | | 2.46 | 0.70 | |
| TLR8 | 2.71 | 1.90 | 3.33 | 2.22 | 2.89 |
| TLR9 | | | | | |
| TNF | 4.37 | 4.16 | 2.80 | 2.47 | 4.38 |
| TNFAIP3 | -0.12 | 0.45 | 0.12 | -0.07 | 1.38 |
| TNFSF14 | | | | | |
| TOLLIP | -0.39 | -0.11 | -0.21 | -0.18 | 0.49 |
| TRADD | -0.77 | -0.13 | 0.09 | 0.83 | 0.71 |
| TRAF2 | | | 0.02 | -0.45 | |
| TREM2 | | | | | |
| TSLP | | | | | |
| TWIST2 | | | | | |
| TYROBP | 7.29 | 6.24 | 3.79 | 4.67 | 6.89 |

$Log_2$ Fold Change Immune/non-immune (scale: 8, 6, 4, 2, 0, -2, -4, -6, -8)
